# Rarity: Discovering rare cell populations from single-cell imaging data

**DOI:** 10.1101/2022.07.15.500256

**Authors:** Kaspar Märtens, Michele Bortolomeazzi, Lucia Montorsi, Jo Spencer, Francesca Ciccarelli, Christopher Yau

## Abstract

**Background:** Cell type identification plays an important role in the analysis and interpretation of single-cell data and can be carried out via supervised or unsupervised clustering approaches. Supervised methods are best suited where we can list all cell types and their respective marker genes *a priori*. While unsupervised clustering algorithms look for groups of cells with similar expression properties. This property permits the identification of both known and unknown cell populations, making unsupervised methods suitable for discovery.. Success is dependent on the relative strength of the expression signature of each group as well as the number of cells. Rare cell types therefore present a particular challenge that are magnified when they are defined by differentially expressing a small number of genes. Typical unsupervised approaches fail to identify such rare subpopulations, and these cells tend to be absorbed into more prevalent cell types.

**Results:** In order to balance these competing demands, we have developed a novel statistical framework for unsupervised clustering, named *Rarity,* that enables the discovery process for rare cell types to be more robust, consistent and interpretable. We achieve this by devising a novel clustering method based on a Bayesian latent variable model in which we assign cells to inferred latent binary on/off expression profiles. This lets us achieve increased sensitivity to rare cell populations while also allowing us to control and interpret potential false positive discoveries.

**Conclusions:** We systematically study the challenges associated with rare cell type identification and demonstrate the utility of Rarity on various IMC data sets.

## Background

High-dimensional molecular analysis of single cells with highly-multiplex imaging allows the simultaneous measurement of the expression of multiple proteins while retaining information about their spatial origin within the tissue section. Technologies such as imaging mass cytometry (IMC) (Giesen et al. 2014) and multiplexed ion beam imaging (MIBI) (Angelo et al. 2014) use antibodies conjugated with heavy metals to stain tissues, which is followed by laser ablation and mass spectrometry to quantify expression of around 40 pre-determined molecular markers. Immunofluorescence-based microscopy such as multiplexed immunofluorescence (Gerdes et al. 2013) and cyclic immunofluorescence (CyCIF) (Lin et al. 2018), allow the multiplexed detection of proteins using standard microscopy.

A high-throughput single cell analysis using such technologies can therefore lead to molecular profiles of tens of thousands of cells. For instance, (Damond et al. 2019) used IMC to analyse and characterise the pathogenesis and progression of Type 1 Diabetes in human patients. While (M.Bortolomeazzi, Keddar, and Montorsi 2020) integrated multi-regional whole-exome, RNA and T-cell receptor sequencing as well as IMC to examine the tumor microenvironment of hypermutated colorectal cancers in response to anti-PD1 immunotherapy. MIBI-TOF (Keren et al. 2018) was used to profile 36 immune-related proteins (including PD1, PD-L1, and IDO) in 41 triple-negative breast cancer patients to reveal mixed and compartmentalized tumors that coincided with cell type and location specific expression of key markers.

A standard step in single cell analysis is cell type identification and classification in which cells (data points) are sorted into phenotypically distinct groups (clusters). This can be accomplished via *supervised* or *unsupervised* clustering approaches. Supervised clustering algorithms (Abdelaal et al. 2019; Geuenich et al. 2021) require *a priori* information in terms of either pre-annotated cells or cell type profiles so that cells can be suitably labelled and assigned to one of these known cell types. In contrast, unsupervised clustering algorithms (Levine et al. 2015; Van Gassen et al. 2015) look for groups of cells exhibiting “similar” expression profiles, assuming that this latent clustering structure in the data will coincide with cell types, and require no *a priori* information. Therefore unsupervised approaches preclude the need to pre-specify the precise number of clusters and their nature in advance but require more post-hoc effort to annotate the clusters upon discovery. Unsupervised approaches are best suited for *discovery* of novel cell phenotypes, while supervised methods are ideal when the landscape of cell types is already well-characterised. A number of computational packages are now available to wraparound and simplify the use of such analysis (Eling et al. 2020; Opzoomer et al. 2021).

One particular challenge is the identification of novel *rare* cell populations. A review of eighteen clustering methods in (Weber and Robinson 2016), including *FlowSOM* (Van Gassen et al. 2015) and *PhenoGraph* (Levine et al. 2015), across six high-dimensional single cell flow and mass cytometry data demonstrated a wide variation in performance in rare cell population detection. This is unsurprising since the clustering approaches were typically unsupervised algorithms designed to identify subgroups in the data with no explicit criteria adopted to specify that some subgroups may be small. Evaluation in the review was restricted to the ability to recover one or two predefined rare cell populations.

Existing unsupervised methods are often limited in their ability to identify such populations whose presence can be occluded by more prevalent cell populations. Unsupervised algorithms struggle to identify rare clusters as their success is dependent on the relative strength of the expression signature of each group compared to others and importantly the number of cells in each group. This is in contrast to supervised clustering algorithms that identify cells exhibiting a particular known pattern of expression. The latter process does not in principle depend on sample size, so rare, but known, cell types can be more readily identified. Nevertheless, accuracy may be variable across cell types, depending on how discriminative and specific the cell markers are) and does require that the cell type of interest and its characteristics are fully known. The key feature that underpins our proposed methodology, to be presented later, is to combine the marker-based approach of supervised algorithms with the ability to discover novel groups in unsupervised algorithms.

A major obstacle with developing methods to detect rare and unknown cell types is the need to balance the competing forces of postulating novel clusters and the production of false positives. While there must be an acceptance that increased sensitivity for rare cell types will increase false positives, this must be managed in order to minimise downstream complications in interpretation and simplified exploration of putative clusters for analysts and experimentalists alike. Further, unsupervised algorithms often contain adjustable parameters to vary performance and these can be used to produce outputs with more or fewer clusters and therefore potentially more rare clusters. However, the interpretation of results arising can be problematic. The overall clustering structure can be radically different across different parameterisations and therefore changes in algorithmic parameters do not lead to “smooth” changes in clustering patterns and can in fact lead to quite contradictory clusters. This instability is also noted in (Weber and Robinson 2016).

As a result, we have developed *Rarity,* a hybrid semi-supervised framework for cell type identification that has been specifically developed to be sensitive to rare subpopulations. We demonstrate that Rarity is able to identify putative rare cell populations that existing clustering methods do not identify and cannot be visualised with high-dimensional visualisation techniques such as t-SNE (van der Maaten 2008) and UMAP (McInnes, Healy, and Melville 2018; Becht et al. 2018). Further, through the use of a binary latent feature model, we illustrate how Rarity assigns a simple and interpretable binary marker signature to each cluster making post-hoc examination, filtering and verification of clusters substantially easier.

Figure 1 illustrates the aforementioned challenges using a synthetic dataset (see Methods for simulation details) containing three *known* and two *unknown* cell populations — the former (cell types A–C) are present at a high prevalence (49%, 33% and 12% of cells respectively) whereas the latter (cell types D–E) are rare (with prevalence below 1%). The zoom-in provided in Figure 1A highlights how the UMAP dimensionality reduction has not recognised cell type D as a recognisably distinct cluster, instead it is mixed with the more prevalent cell types A and B. When applying a supervised model *Astir* (Geuenich et al. 2021) after specifying the marker genes for the known cell types A-C, Astir assigns both rare cell types to an “Other” category (Figure 1B) but does not provide functionality for any further analysis of this “Other”category. When applying an unsupervised model *Phenograph* (*Levine et al. 2015*) (Figure 1C), we end up with 15 clusters and the true rare cell type D is still split across four clusters. In contrast, our proposed approach *Rarity* (Figure 1D) allows us to distinguish between known and unknown cell types, and interpret the clusters in terms of their marker signatures.

**Figure 1.**
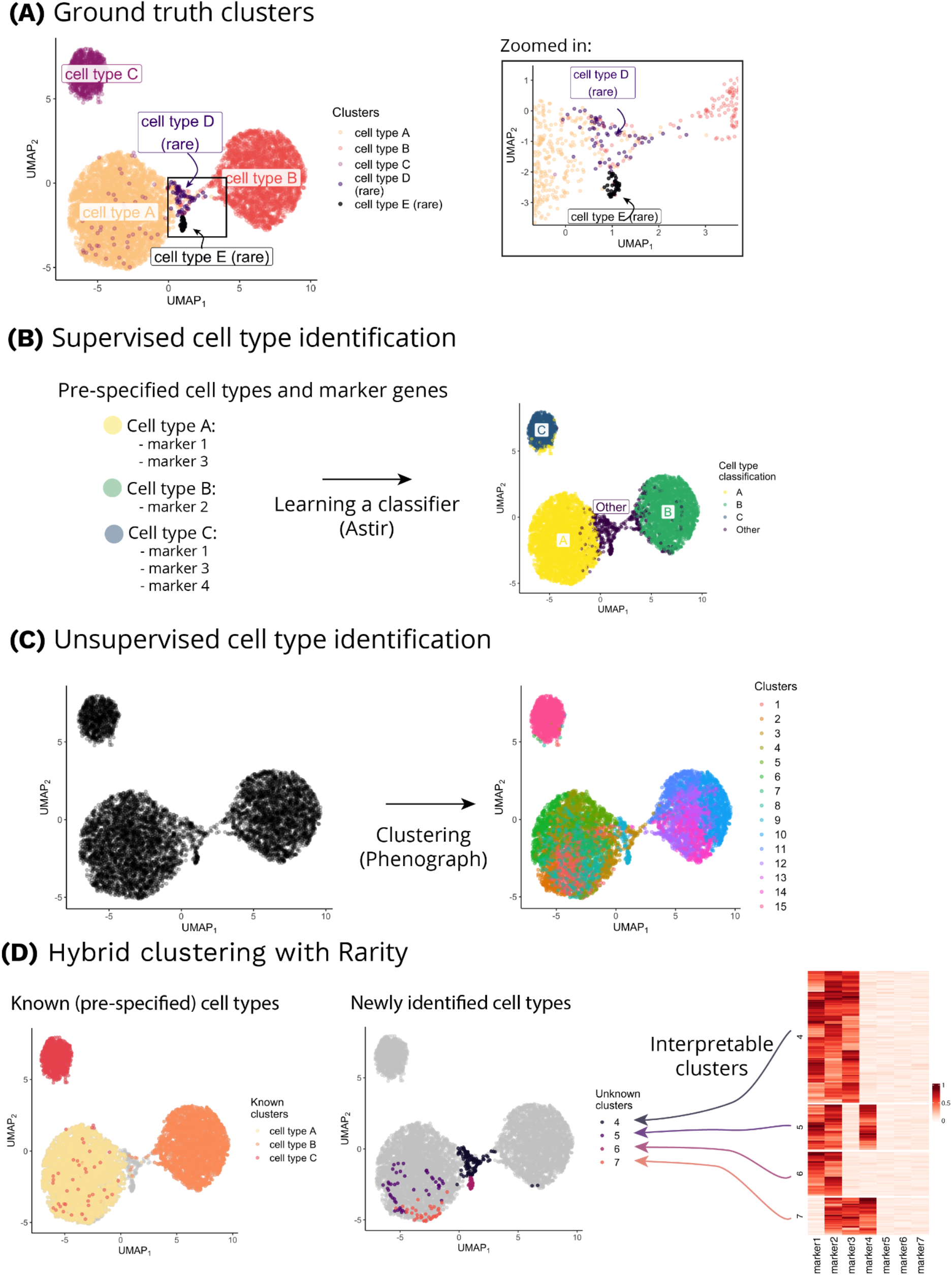
Comparison of supervised and unsupervised workflows for cell type identification on a synthetic dataset highlights their drawbacks motivating our hybrid modelling framework. (A) The synthetic dataset comprises five different cell populations (cell types A-E), two of which are rare and unknown to us *a priori* (cell types D and E). The UMAP plot is coloured by ground truth cell type labels. The zoomed-in panel shows a close-up of the two rare cell types (note that the UMAP visualisation has failed to recognise cell type D as a distinct cluster). (B) In the supervised case (here shown for *Astir(Geuenich et al. 2021)*), our capabilities to detect cell types are limited to the pre-specified cell types and their characteristic markers. The rare cell types are either merged with other known cell types or assigned to a separate “Other” cluster. (C) In the unsupervised case, the workflow involves running a clustering algorithm (here shown for *Phenograph*), followed by manual inspection of marker genes in order to label (and potentially merge) the inferred clusters. (D) The proposed hybrid approach *Rarity* can help in identifying both pre-specified cell types (as shown on the left UMAP) as well as rare novel cell populations (as shown on the right UMAP). The identified clusters are interpretable in terms of their differential expression profile—the identified rare clusters are shown in the heatmap.

Our contributions are threefold. First, we systematically study and highlight limitations of existing clustering approaches for rare cell type identification in various simulation studies, including downsampling experiments on real IMC data. Second, we introduce *Rarity* and show how it successfully addresses some of the limitations of existing methods. Third, we present an analysis of a colon mucosa IMC dataset, showing how *Rarity* leads to the identification of rare, double negative, CD4- and CD8-T cells.

## Results

### Rarity: A hybrid clustering framework for detecting rare cell populations

Rarity combines the benefits of supervised and unsupervised approaches to cell type identification in a hybrid probabilistic framework. It was designed to be sensitive to rare subpopulations, including those which differ from other cell types in the expression of even a single marker. To achieve this level of sensitivity without sacrificing interpretability, we have built Rarity upon a statistical modelling assumption that the continuous marker expression values associated with each cell have an underlying binary on/off state. We model these unobserved on/off states as binary latent variables. We then associate every unique binary signature with a cluster label. As a result, every cell is mapped to a latent cluster that is associated with a particular binary gene expression signature (Figure 2A). Known cell types can therefore be specified by *a priori* specifying the appropriate binary expression pattern.

**Figure 2.**
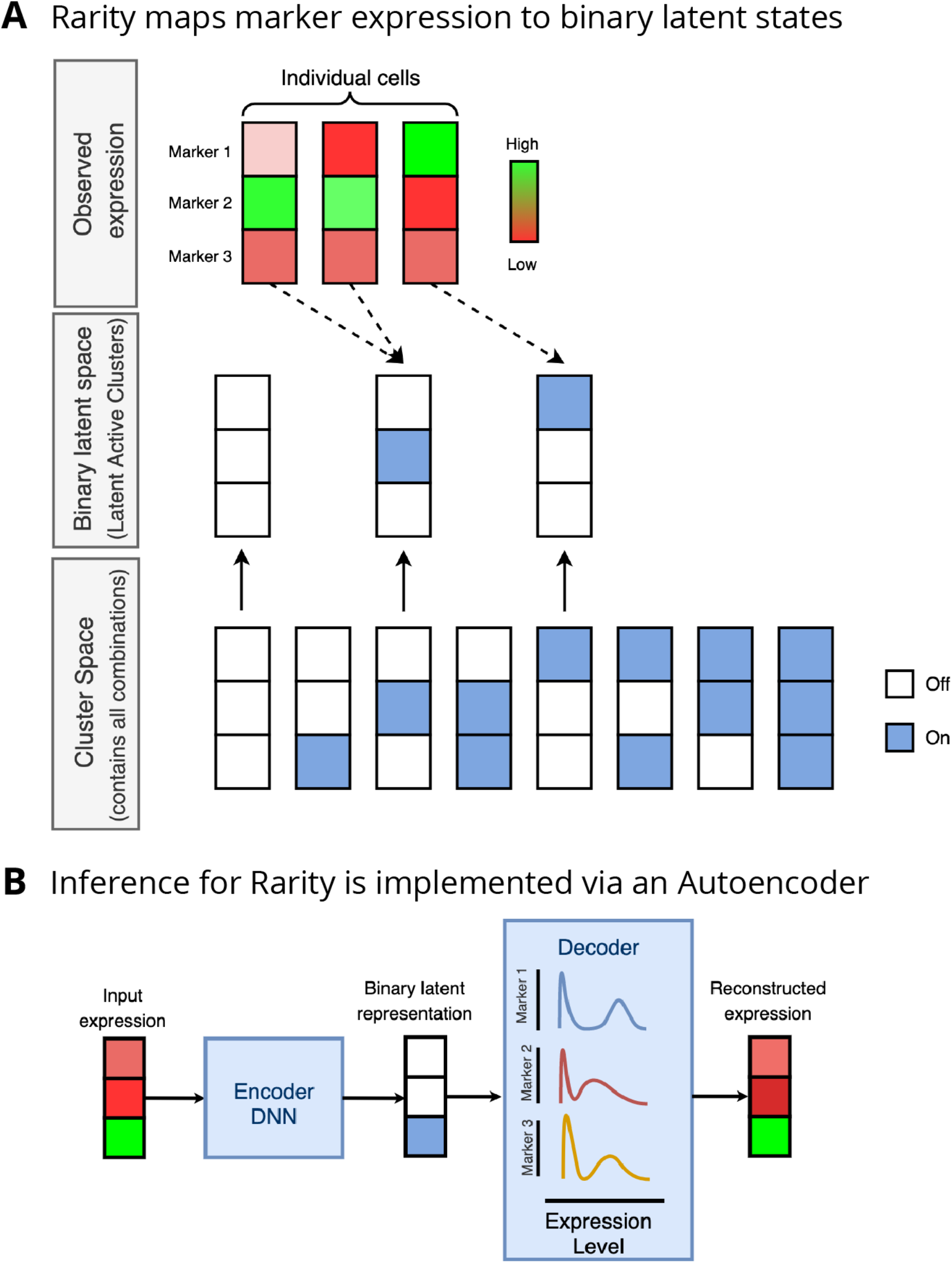
Overview of Rarity model specification and implementation. (A) Rarity projects single cell marker intensity vectors (illustrated for three markers and three cells, top panel) onto binary expression signatures (in the middle panel). Note that the *active* latent signatures cover only a subset of all possible binary combinations in the cluster space (in the bottom panel). (B) Tractable inference for Rarity is implemented as a structured autoencoder, where an encoder neural network is used in combination with continuous relaxations to project expression vectors to binary expression signatures.

Modelling the underlying binary states comes with a computational challenge, since with *M* markers there are a maximum of 2^M^ possible binary expression signatures. To identify which of these a cell belongs to, we would then need to consider 2^M^ potential cell types for every cell. We transform this intractable combinatorial complexity into a tractable computation by reformulating the binary inference problem by using continuous relaxations (Maddison, Mnih, and Teh 2017). That is, Rarity performs inference for the binary signatures by mapping single cell data vectors onto continuous relaxations of binary encodings. This allows us to benefit from the use of modern differentiable computing frameworks such as PyTorch (Paszke et al. 2019) and enables training of the model without the need to explicitly consider all possible binary configurations (Figure 2).

Rarity is implemented in an autoencoder framework (Figure 2B). As a result, our implementation also scales favourably to a large number of cells. We employ *inference amortisation*, an approximate inference technique which introduces an encoder neural network as a form of parameter sharing (Kingma and Welling 2014; Rezende, Mohamed, and Wierstra 2014) across cells. As a result, even though the binary expression signatures are inferred for every cell individually, the number of learnable parameters is fixed and does not grow with the number of cells in our data. Moreover, having a trained model, we can simply employ it on new cells without additional training.

Code and use case examples for Rarity are available from a Github repository https://github.com/kasparmartens/rarity.

### Self-consistency clustering experimental procedures and metrics

To investigate the sensitivity of existing IMC clustering methods for detecting cell types present at various prevalence levels in a controlled setting, we need to resort to either synthetic or semi-synthetic simulation experiments. We first outline in detail the strategy for these simulation experiments and the corresponding benchmarking metrics which we will adopt throughout.

Our experiments involve running a clustering algorithm on a simulated or real data set to derive clusters corresponding to cell types. We will then downsample one of the identified clusters to create a new dataset with fewer cells of that type and re-cluster using the same method. The goal of this experiment is to measure how similar (or different) the inferred clusters from the downsampled data set will be from the original one. A highly desirable property of a clustering method is *self-consistency,* i.e. the ability to allocate cells in the same cluster regardless of the prevalence of each cluster (such as after downsampling).

To quantify the quality of the inferred clustering, we adopt two metrics that capture different aspects of self-consistency. First we measure *homogeneity,* i.e. the property of the inferred clusters to contain only a distinct cell type. Second, we measure *completeness,* i.e. the property of grouping all cells of a particular type in only one cluster (Rosenberg and Hirschberg 2007). Since our focus lies in the identification of rare cell types, in our downsampling experiments we measure these two scores conditional on the downsampled cluster. We refer to these conditional scores as *conditional homogeneity* and *conditional completeness,* both with values between 0 and 1, higher scores being better (see Methods for more details). To further summarise these two scores with a single summary statistic, we use the harmonic average of the two, the conditional V-measure (Rosenberg and Hirschberg 2007).

The behaviour of these metrics is illustrated in Figure 3 for four different clustering scenarios (Clustering 1-4). Suppose a clustering method identifies three cell types (red, yellow and blue) from the original data. We then reduce the number of blue cells to create a downsampled data set. In the first scenario (Clustering 1), the method identifies two clusters (green/purple) from the downsampled dataset. The completeness is high as all cells in each of the original clusters map to only one of the new clusters (orange to green, red to purple). However, homogeneity is low as the purple cluster contains both blue and red cells. In contrast, in the second scenario (Clustering 2), the method finds four clusters. The result is homogeneous as all the original orange cells are in their own new cluster (green) and all red cells are in another new cluster (purple). However, the blue cells are split into new blue and yellow clusters so completeness is low. Alternatively, the method could discover three clusters (Clustering 3) but under a different configuration of labelling. While all orange cells map to the green cluster, the blue and red cells map to two new clusters (purple/yellow) which partition across the original blue and red clusters. Thus homogeneity and completeness scores are low. Only in the perfect scenario (Clustering 4) where the method perfectly reclassifies on the downsampled data would all metrics have value 1.0.

**Figure 3.**
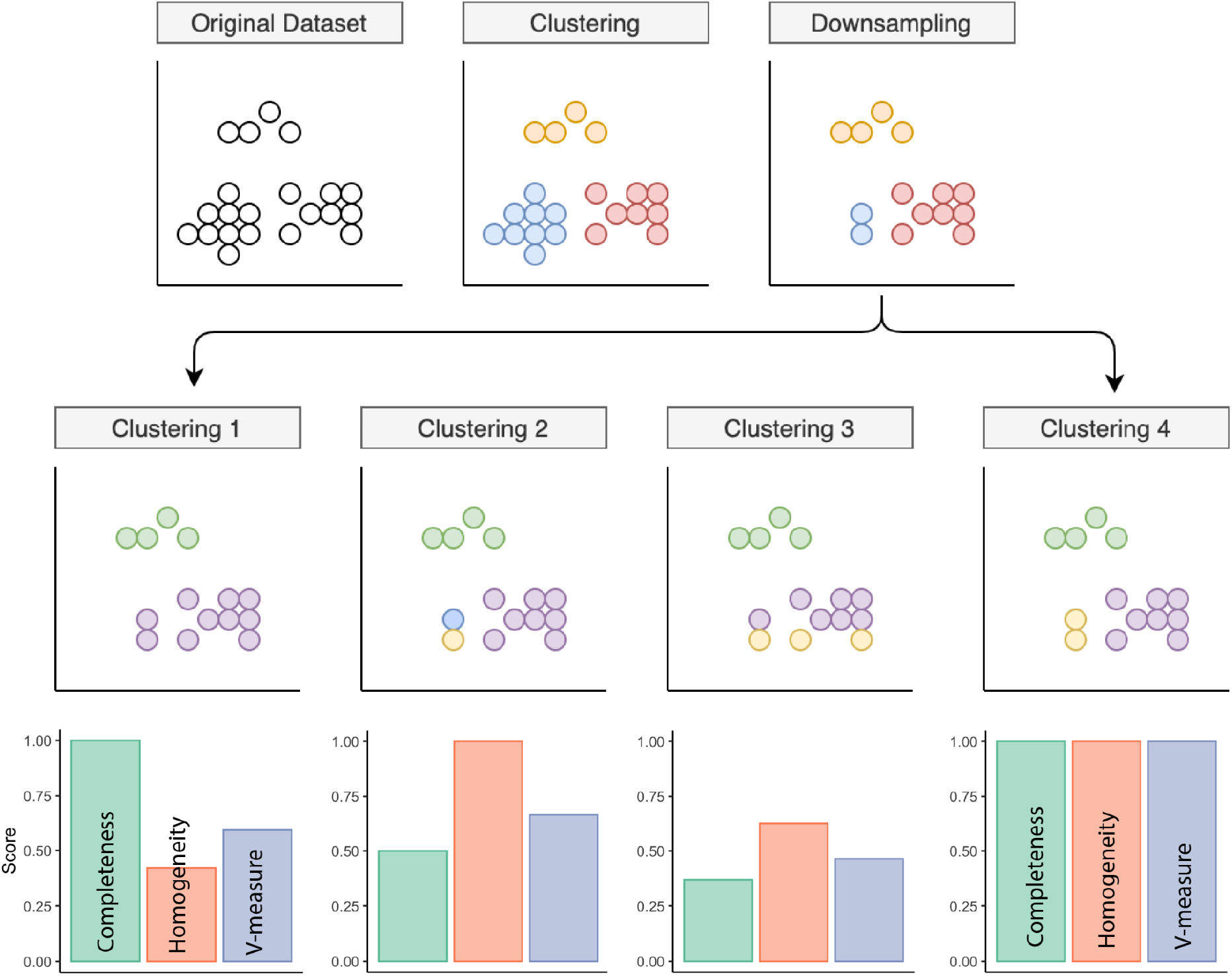
Downsampling simulation experiments and benchmark metrics to quantify self-consistency. Schematic showing how the downsampling experiments were conducted, and how different outcome scenarios affect the clustering performance metrics.

### Existing unsupervised clustering methods fail to reliably detect rare cell populations

Given this experimental setup, we then sought to examine how effective existing commonly used unsupervised clustering methods (PhenoGraph and FlowSOM) were at detecting rare cell populations and compared the performance of Rarity relative to this. We devised a simulation experiment in which we generated artificial data sets consisting of five cell types (Figure 4A) with three common (cell types A, B, C) and two rare (cell types D, E) where the latter were downsampled from an initial 5% of the cell population, to 1% and then 0.5%. Using PhenoGraph (with the number of nearest neighbours set to the default value 30), we observed that common cell populations had the tendency to be fragmented into multiple clusters by the algorithm while the ability to detect rare populations diminished as the population size decreased (Figure 4B). In contrast, with Rarity, common and rare cell populations were more reliably and consistently identified even with the decreasing rare cell population size (Figure 4C). For comparison, we also assigned cell types with a supervised method, Astir, providing the marker information for all five cell types (Supplementary Figure S2). Using our previously introduced clustering metrics, Rarity (Figure 4D) showed consistently superior performance compared to both PhenoGraph and Astir even though the latter is given the cell type profiles. In particular, while PhenoGraph maintains relatively high completeness, signifying that it tends to merge rare cell types in one cluster, homogeneity is low, as more than one cell type can be mapped to the same cluster. This illustrates the need for dual metrics to understand the complexities of interpreting clustering output where the entire cluster structure may alter under different conditions.

**Figure 4.**
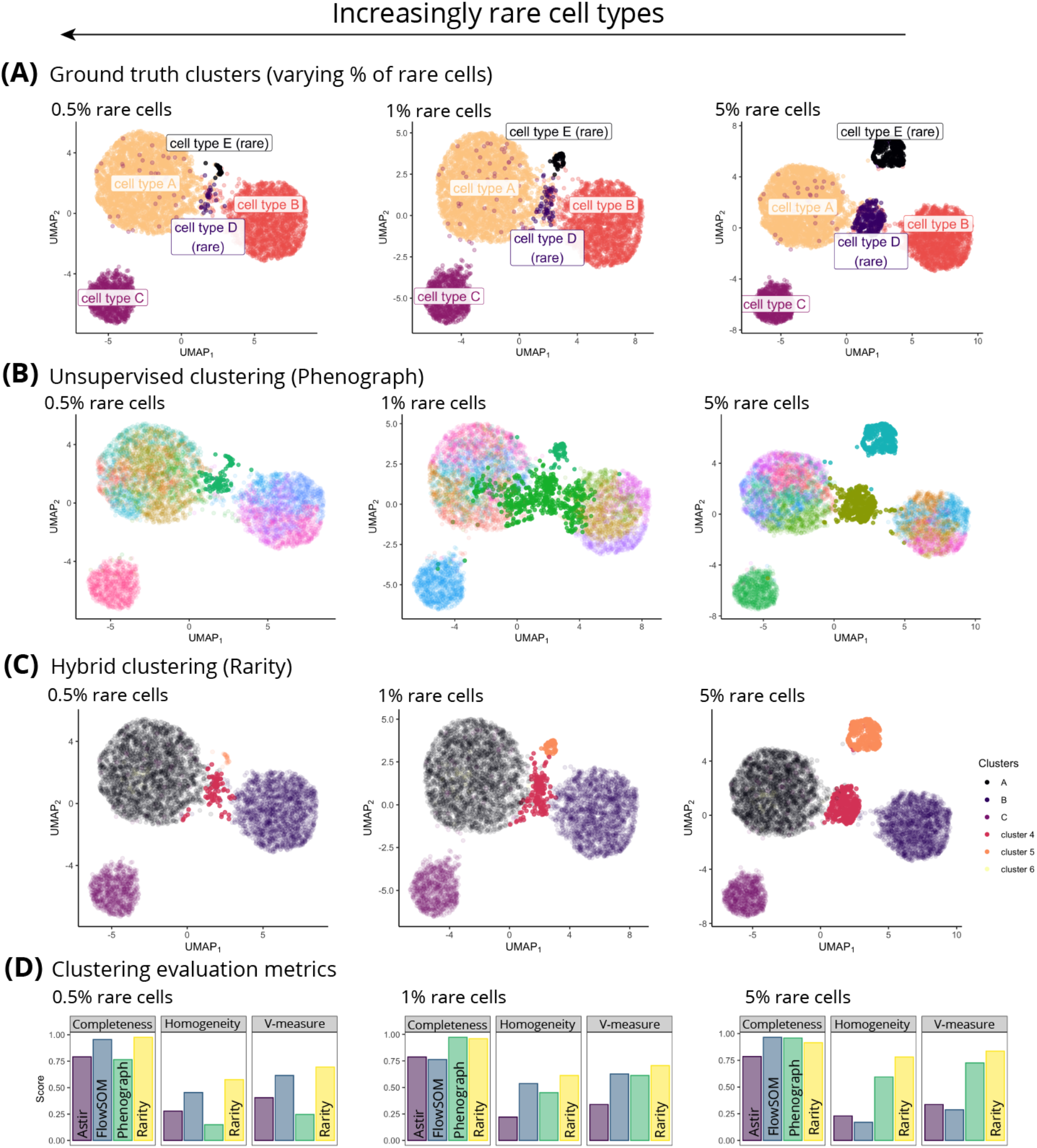
The less prevalent a cell type is, the more challenging it is to be reliably identified with unsupervised clustering methods. **A.** Similarly to Figure 1, ground truth data contains three common and two rare cell types. The three panels represent scenarios with varying extent of rarity: 0.5%, 1%, and 5% prevalence of rare cell types. B. Unsupervised clustering with Phenograph (clusters shown in colour, highlighting the clusters which have the largest overlap with rare cell types) works well when the rare clusters are present at the 5% fraction, however it has failed to identify one of the rare subpopulations at 1% presence, and has only partially grouped the two rare cell types together at 0.5% prevalence. C. Hybrid clustering with Rarity (clusters shown in colour) has correctly identified the two rare groups in all three scenarios. D. To quantify clustering performance for (i) supervised (Astir), (ii) unsupervised (Phenograph) and (iii) hybrid (Rarity) methods, we display the conditional completeness, homogeneity, and V-measure scores (higher is better).

### Breast cancer IMC data

We next considered a breast cancer IMC dataset (Jackson et al. 2020) and conducted an analysis using PhenoGraph and FlowSOM (Van Gassen et al. 2015) specifically looking at their ability to identify novel rare sub-populations before considering the utility of Rarity. Both PhenoGraph and FlowSOM possess algorithmic parameters which can be modified to enable these methods to produce different numbers of output clusters - including potentially those corresponding to putative small subpopulations. Figure 5 shows how the number of clusters reported by PhenoGraph (Figure 5A) and FlowSOM (Figure 5B) varies with these parameters.

**Figure 5.**
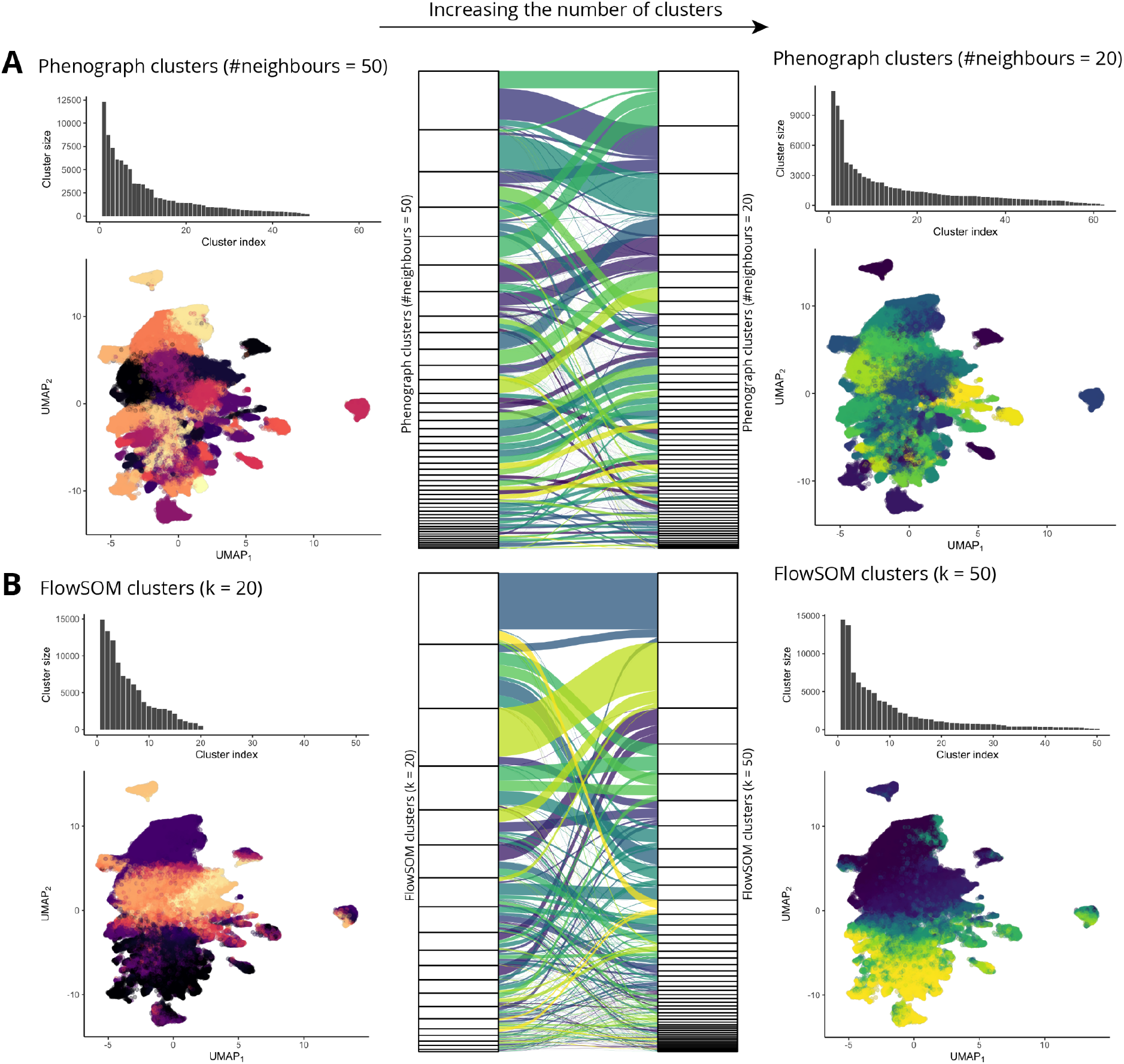
Unsupervised clustering algorithms have hyperparameters which let us either directly (FlowSOM) or indirectly (Phenograph) control the number of clusters, but clusters obtained under different hyperparameter configurations are not consistent to one another. (A) When we decrease the number of nearest neighbours in Phenograph from 50 (on the left) to 20 (on the right), we effectively increase the number of clusters, however the mapping between the two sets of clusters (shown in the alluvial diagram in the middle) is highly complex and indicates inconsistency, as many clusters are both split and merged. (B) Similarly for FlowSOM, when we increase the number of clusters from 20 (on the left) to 50 (on the right), we observe both cluster splitting as well as merging.

When the PhenoGraph hyperparameter corresponding to the number of neighbours is reduced from 50 to 20 (Figure 5A), the number of clusters discovered increases from 48 to 62 and there is an increase in the number of clusters which represent less than 1% of the total population from 24 to 35. In contrast, in FlowSOM we can explicitly control the number of clusters reported and we demonstrate this when increasing this number from 20 to 50 (Figure 5B). This change led to an increase in the number of clusters which represent less than 1% of the total population from 3 to 30.

We next sought to understand how the underlying cluster structure alters as cluster number changes. Figure 5 illustrates how individual cell cluster assignments vary with output cluster number for FlowSOM and PhenoGraph. As cluster number changes, there are substantial cluster structure alterations with some clusters merging and splitting as more clusters are returned. For PhenoGraph, there was an average of 11 parents from the original clustering for each of the 62 clusters, while each of the 50 FlowSOM clusters had 7 parents. This indicates that for both methods, increases in the number of clusters did not increase sensitivity and the detection of small rare populations, instead it led to fundamental changes in clustering structure.

When examining how particular clusters are split in terms of marker expression levels, Figure 6A shows an illustrative example focusing on epithelial luminal cells, where PhenoGraph has generated two daughter clusters from a parent cluster. Each daughter cluster differs from the other only via a subtle change in the expression of two markers. For a similar group of epithelial luminal cells, FlowSOM partitions the cluster into two daughter clusters with entirely different expression signatures (Figure 6B). These outputs are challenging to interpret as cluster structure and properties fundamentally alter as greater sensitivity is encouraged.

**Figure 6.**
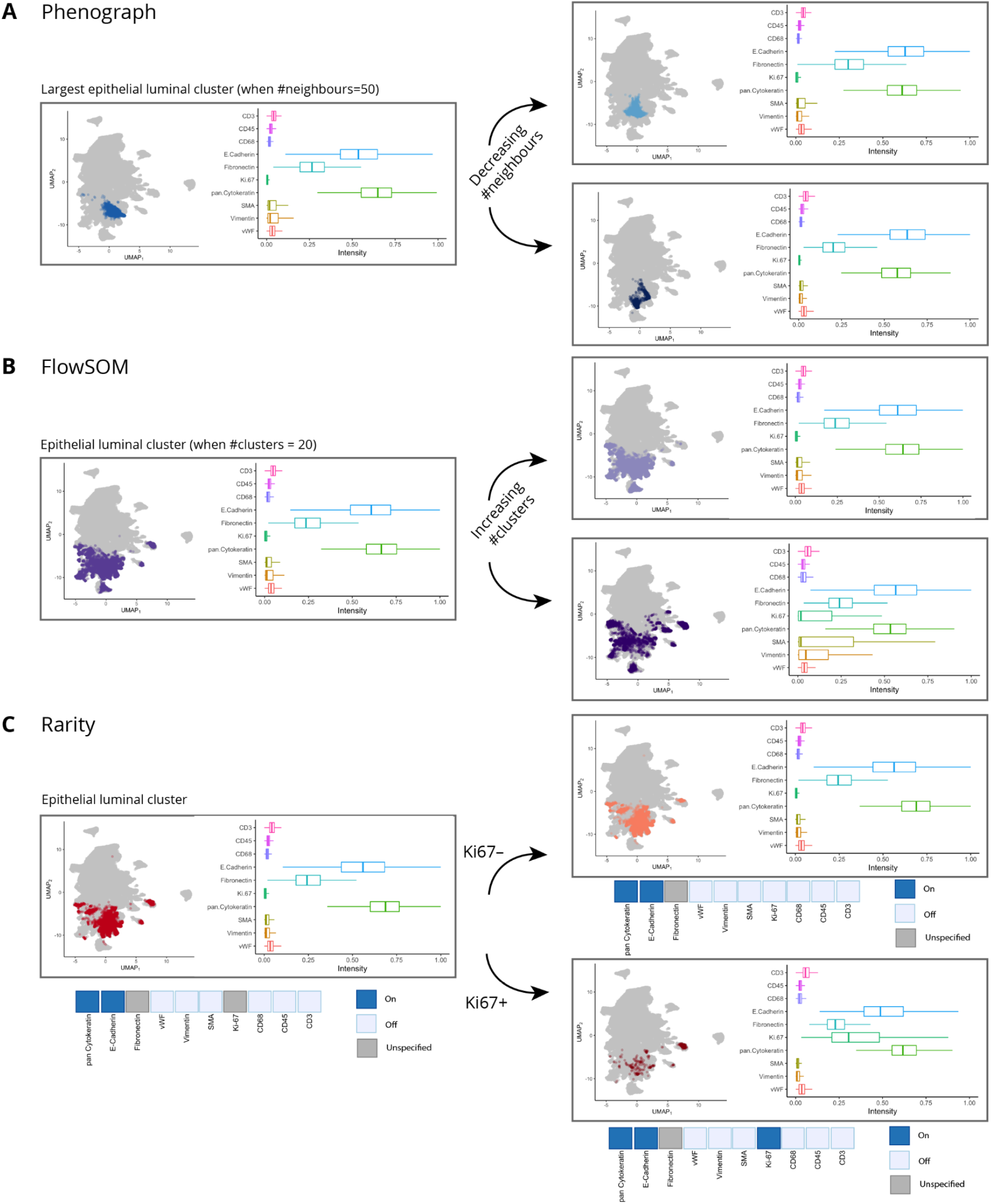
Increasing the number of clusters in unsupervised clustering methods does not typically help with identifying subclusters with distinct expression signatures. In all panels (A-C), we show the largest epithelial luminal cluster identified by (A) Phenograph, (B) FlowSOM, (C) Rarity, both on the UMAP plot as well as on the corresponding expression values boxplot. When increasing granularity (the clusters shown in the right column), the subpopulations identified by Phenograph are both extremely similar. For FlowSOM, a subpopulation expressing moderate levels of various markers (Vimentin, SMA, Ki67) emerges. In contrast, Rarity is the only method, where increase in granularity is directly interpretable in terms of expressing/not expressing a marker. The split illustrates how Ki67+ subpopulation of epithelial luminal cells can be identified with Rarity.

The hybrid approach of Rarity leads to a different approach to cluster interpretation. Since IMC profiles are mapped to latent binary expression vectors, we can select cells that match a particular expression pattern to determine clusters. Here, we were able to select a group of epithelial luminal cells by interrogating all cells that were mapped by Rarity to any latent binary vectors that express E-Cadherin and pan-Cytokeratin, but do not express e.g. CD45 or CD3 (Figure 6C, full list of markers provided in Supplementary Table 3). We can then examine this set of cells and ask which cells among these do and which do not express Ki-67 by creating a more refined query for binary expression vectors, now additionally accounting for the presence and absence of Ki-67 expression.

With Rarity, each cell is mapped onto a cluster with a clear latent binary expression pattern, where each cluster (by definition) must at least differ by the expression of one marker (Figure 6C, Figure 7) and leads to a natural hierarchy of clustering assignments.

**Figure 7.**
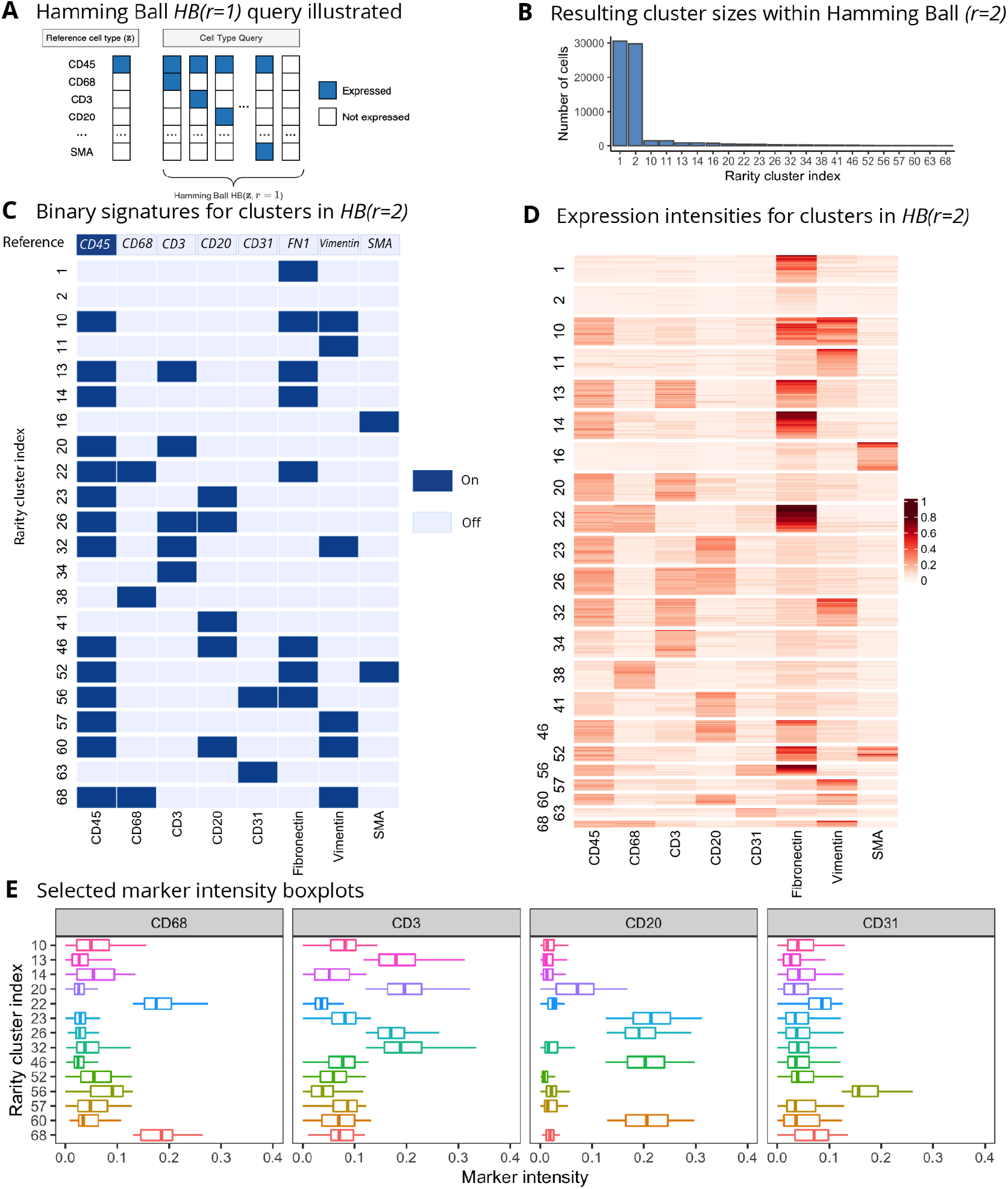
Rarity finds interpretable substructure within cells identified as “Other” by Astir on the breast cancer data. (A) Hamming Ball query with radius *r=1* illustrated: given a reference signature that expresses the CD45 marker, the query returns a set of binary signatures that differ from this reference at one *(r=1)* or two *(r=2)* markers. Panels (B-E) show the results for a Hamming Ball query with the same reference but now *r=2*. (B) The number of cells in each cluster within the Hamming Ball. (C) The binary signatures for all clusters within the Hamming Ball, and (D) the corresponding observed intensities heatmap (for visual aid, we display a subset of the two largest clusters on the heatmap). (E) Selected marker intensity boxplots (CD68, CD3, CD20, CD31) for the identified clusters (shown for all clusters that express CD45) are in concordance with the binary profiles shown in (C) and highlight how every cluster has a unique interpretation in terms of differential expression signatures.

Next we demonstrate further how Rarity can help us identify a more granular clustering which can give insights into finding putative rare cell subpopulations. One way to explore the subpopulations identified by Rarity is via what we call the “Hamming Ball query” (Figure 7A) where we search for binary expression signatures that deviate from the signature of a reference cell type no more than a given number of markers (i.e. their Hamming distance from the reference signature does not exceed a given radius). For example, Figure 7A illustrates a query where the reference signature corresponds to cells that express the CD45 marker. This query would let us identify various immune cells, for example the Hamming Ball with radius *r=1* would contain macrophages, i.e. a signature where both CD45 and CD68 markers are expressed.

This functionality shows how Rarity can complement the cell type identification with a supervised method such as Astir (Geuenich et al. 2021) where all the cell types of interest have to be pre-specified *a priori*. In the case of Astir, there is an additional “Other” cluster that will consist of a mix of unrecognised cell types. Figure 7 shows how Rarity has found substructure within this “Other” group of cells. Specifically, we aim to distinguish between different classes of stromal cells: immune cells, endothelial cells, smooth muscle and fibroblasts. We have conducted analysis starting from the reference signature with CD45 as the main immune cell marker (binary expression signature shown in the top row of panel C), and considering all cell type signatures within the Hamming Ball with radius *r*=2. For example, the first (and largest) cluster does not express the CD45 marker but does express Vimentin, suggesting that it corresponds to a set of cells from the connective tissue. Clusters that co-express both CD45 and Fibronectin/Vimentin (i.e. clusters 10, 13, 14, 22, 32 etc) are likely to be immune cells that are located within the stroma, e.g. cluster number 13 would correspond to such T cells. Zooming in to clusters which express CD68 (i.e. clusters 22, 38, 68) helps us identify macrophages. Rarity has also identified a group of vascular cells (clusters 56 and 63 which express the CD31 marker). The heatmap (Figure 7D) and boxplots (Figure 7E) confirm that indeed the inferred binary signatures are indicative of the actual intensity levels, and thus aiding interpretability.

### Downsampling experiments highlight how most clustering methods are inconsistent

We next performed a downsampling experiment using the same breast cancer IMC data to further characterise the rare cell detection capabilities of each method. We applied FlowSOM, PhenoGraph, Louvain clustering implemented within Seurat v3 (Stuart et al. 2019), as well as the supervised method Astir together with Rarity to the data. Each method identified different clustering configurations and hence different cell type numbers. However, from the output of each method, we identified the clusters most likely to correspond to epithelial or T cells (using the expression of E-Cadherin and pan-Cytokeratin for epithelial, and CD45, CD3 for T cells). For Astir, these markers were used to pre-define the cell signature to be identified as input. We then created downsampled datasets. First, downsampling epithelial cells in one experiment and T cells in the next. These cell types were reduced from the original numbers to 1000, 250 and 100 cells respectively (Figure 8). The clustering methods were then reapplied to the downsampled data to determine if the clustering labels from the full data set are recapitulated. Thus, these experiments do not rely on any “ground truth” cell type labels, they simply indicate how consistent every method on its own is.

**Figure 8.**
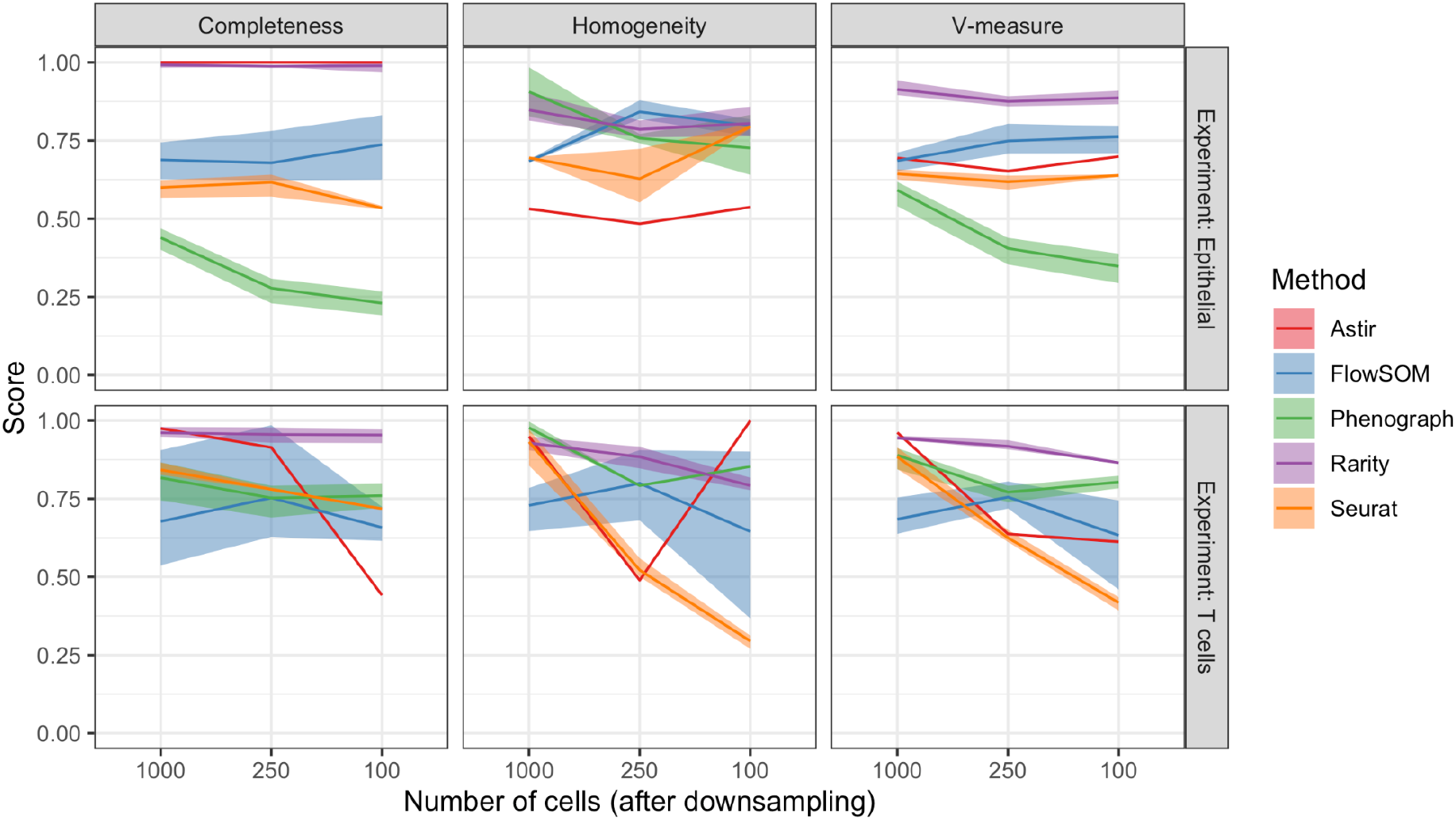
Downsampling simulation experiments quantifying self-consistency. Clustering scores (conditional completeness, homogeneity, and V-measure) when downsampling the largest Epithelial (top row) or T-cell (bottom row) cluster identified by the respective methods (Astir, FlowSOM, Phenograph, Rarity, Seurat), varying the number of cells retained after downsampling from 1000 to 250 to 100 (x-axis). The scores quantity the goodness of the clustering w.r.t. the respective methods run on the full dataset, thus quantifying self-consistency.

Our experiments showed that Rarity showed stable and superior clustering performance to that of other unsupervised approaches (Figure 8). We observed that the sensitivity of unsupervised methods did not simply decline with decreasing numbers of cells, but that the performance characteristics were more complex and there was a dependency on cell type. This is due to the fact that clustering output from the same method on original and downsampled data sets often exhibited significantly different clustering output (similarly to the previous synthetic data experiments) while in contrast Rarity’s design makes it less susceptible to this. Intriguingly, while Astir is a supervised method and is given target gene signatures as input, its performance was also not always stable across downsampled datasets. This is due to the fact that certain model parameters are inferred dynamically so its performance is also partially data set dependent in this case with T cells.

### Rarity identifies CD4-negative and CD8-negative T cells in colorectal cancer IMC data

We next examined the utility of Rarity for the discovery of gamma-delta T cells in immunogenomic profiles of normal colon mucosa in patients with multiple or single and familial or sporadic colorectal cancer. IMC data was generated for sixteen samples of non-cancerous colon mucosa were obtained from six individuals who underwent surgical resection of colorectal cancers (see Supplementary Information for full experimental details). The imaging data was originally published as part of (Bortolomeazzi et al. 2022), but now we have additionally made available the processed single-cell data set (Märtens et al. 2022). Gamma-delta T cells were expected to constitute less than 10% of T cells in human colon mucosa (Viney, MacDonald, and Spencer 1990), and their characteristic feature is that they are CD3-positive but both CD4-negative and CD8-negative. However, when using standard supervised and unsupervised methods applied to this data set, no such clusters are identified. This is not surprising in the light of our simulation study in Figure 4, even if those cell types were present. Therefore, we wanted to see if the increased sensitivity in Rarity will identify any potential candidates for such CD4- and CD8-double-negative T cells.

After pre-processing steps (see Supplementary for details), our colon mucosa data set contains a total of 40,364 cells. We used Rarity to classify these cells into B cells, T cells, Macrophages, Dendritic cells, Endothelial, Connective tissue cells, and other cells, as illustrated in Figure 9A, using markers listed in Supplementary Table 4. For selected markers and cell types, Figure 9B shows the corresponding marker intensity boxplots.

**Figure 9.**
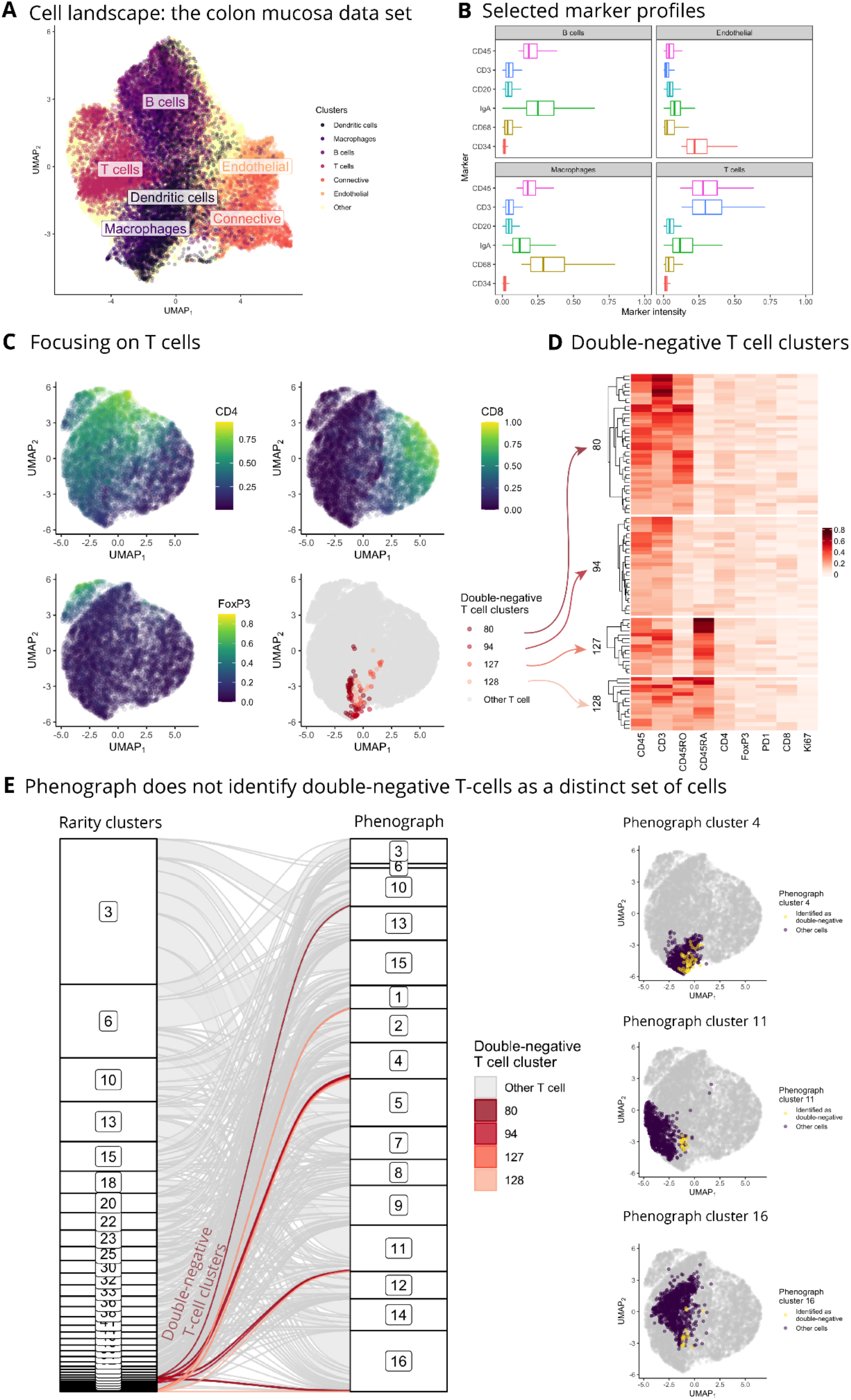
Rarity identifies a unique subpopulation of T-cells that do not express CD4 or CD8 in the colon mucosa data set. (A, B) Cell landscape UMAP plot, where we have highlighted cells identified as B cells, T cells, Macrophages, Dendritic, Endothelial, and Connective tissue cells, together with the corresponding marker intensity boxplots. Next, panels (C-E) focus on analysing T cells only. (C) UMAP plots ofT cells showing expression of CD4, CD8 and FoxP3, and highlighting the location of identified CD4-CD8-T cells (i.e. gamma-delta T cell clusters) by Rarity. (D) Marker expression levels for the identified CD4-CD8-T cell sub-groups (i.e. Rarity clusters 80, 94, 127, 128) show combinatorial co-expression of CD45RA and CD45RO. (E) These double-negative T-cell clusters would not have been identified by unsupervised clustering with Phenograph. In fact, Phenograph has placed these cells into various larger clusters (e.g. clusters number 4, 11, 16 etc) as shown on the alluvial diagram (highlighting the four Rarity clusters) and UMAP plots for selected Phenograph clusters (highlighting the double-negative cells in yellow).

Next, we turned to the analysis of T cells. That is, the following analysis is restricted to the cells identified by Rarity as CD45-positive and CD3-positive. Figure 9C displays the UMAP when re-fitted on T cells only. We can see that CD4 intensity increases along the y-axis, CD8 intensity increases along the x-axis, and the blob in the top left corner corresponds to Regulatory T cells expressing FoxP3. The fourth sub-panel highlights the set of T cells that were identified by Rarity as CD4- and CD8-cells that could potentially be gamma-delta T cells. In fact, Rarity identified four such clusters (clusters number 80, 96, 127, 128). Heatmap in Figure 7D confirms that these clusters have indeed low intensity levels of CD4 and CD8, but it additionally provides insight into why there are four clusters instead of a single one - these clusters are stratified by CD45RO and CD45RA marker intensities indicating memory and naive T cells respectively. In total, the putative gamma-delta T cell clusters lacking CD4 and CD8 comprise 92 cells (which corresponds to 0.2% of all cells and 1.0% of all T cells).

We re-emphasise that these clusters were not identified as a distinct set of cells on the UMAP visualisation, a phenomenon that we already observed earlier in Figures 1 and 4. Furthermore, this set of cells would not have been discovered by an unsupervised method like Phenograph. Figure 7E illustrates how cells in these CD4- and CD8-T cell clusters are spread across multiple larger Phenograph clusters - a behaviour consistent with our earlier findings in Figure 4.

We further explored the candidate gamma-delta T cell by mapping these to their spatial locations and samples of origin. We found that the 92 identified cells were uniformly spread across biological samples (15 out of 16, as highlighted in Supplementary Figure S3) with on average 6 cells per sample. This confirms that the cells were not due to an experimental artefact specific to a subset of samples. Figure 10 displays the spatial location of the identified gamma-delta T cells in three samples. These were located in the sub-epithelial areas of diffuse connective tissue of the lamina propria (Kurd and Robey 2014; Hayday and Gibbons 2008) consistent with the known distribution of such cells and their association with intraepithelial sites.

**Figure 10.**
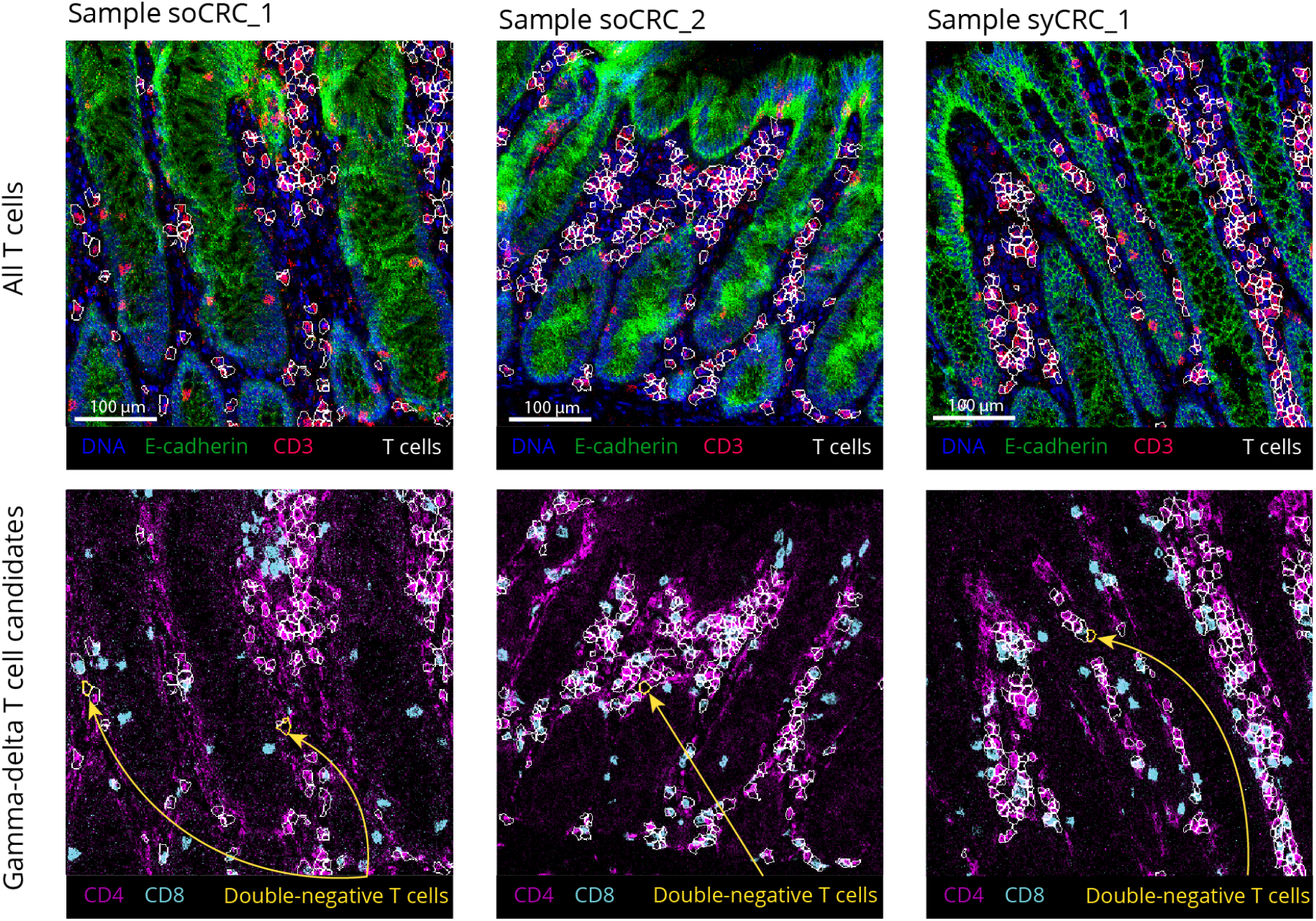
Examples of double-negative T cells identified by Rarity displayed on the original IMC images. For a zoom-in of three samples (soCRC_1, soCRC_2, syCRC_1), the top row shows DNA (blue), E-cadherin (green) and CD3 (red) marker intensities, and highlights T-cell outlines (in white). The bottom row shows CD4 (magenta) and CD8 (cyan) marker intensities, and highlights double-negative T cells (yellow outlines, also indicated with arrows) among all T cells (in white).

## Discussion

Our motivation for the development of Rarity stemmed from the need to identify rare cell types from single-cell data. In this work, we have systematically demonstrated how commonly used clustering methods fail to discover clusters that are rare but yet distinct — our downsampling experiments showed how the task of cell type identification becomes increasingly difficult the less prevalent the cell type becomes. Our simulation experiments also highlight a lack of self-consistency in current unsupervised clustering algorithms.

Given the plethora of literature on clustering methods for cell type detection, *rare* cell type identification has received relatively much less attention. For example, both (Cron et al.11-Jul-2013; Naim et al. 2014) have used a variation of the Gaussian mixture model, with different strategies for choosing the number of clusters. An increased number of clusters will lead to higher sensitivity towards rare subpopulations, however these methods have not been inherently designed to recognise rare groups specifically. As we have demonstrated in the paper, simply increasing the number of clusters does not generally lead to the desired outcome. Instead, increased sensitivity typically leads to breaking existing large clusters down into smaller ones with miniscule differences in their respective expression signatures. Similarly to our findings, the comparative review by (Weber and Robinson 2016) also found that clustering results can be highly variable and e.g. sensitive to bootstrap resampling.

With Rarity we have set out a bespoke approach for delineating these cellular groups. This was based on the idea of transforming the single-cell data into high/low binary expression patterns which induces an implicit clustering of the cells. We demonstrated how this approach led to a robust system for identifying rare cell types and the use of binary expression patterns provides a certain form of guarantee that the clustering output is highly interpretable (i.e. that each cluster must differ from the others by *at least* one molecular feature).

We recognise the limitation of this assumption is that cell types that differ from others only via changes in absolute levels of expression would not be identified by Rarity. To explicitly investigate this limitation, we have modified the synthetic data generative mechanism used in Figures 1 and 4, deviating from the assumption that marker genes are either “on” or “off”, now allowing differential expression to be a continuous-valued quantity Δ (more details in Supplementary Figure S1). Specifically, we focus on relaxing this assumption for the marker gene defining cell type E (expression shown in the boxplot in Figure S1A), where the average intensity of all other cell types is 0.55 but for cell type E it is (0.55-Δ) for varying levels of Δ.

Rarity is most successful in detecting cell type E in scenarios where Δ values are relatively large (0.50 and 0.45), whereas its performance drops significantly for smaller values of Δ (Figure S1C). This demonstrates when Rarity’s binary expression assumption breaks down - indeed for smaller Δ values the average expression intensity for cell type E increasingly deviates from zero, thus becoming challenging to distinguish for Rarity. In contrast, the performance of Phenograph holds more firmly for smaller Δ values. However, this sensitivity is gained through a corresponding increase in the number of false cell types. In this example, Phenograph identifies a total number of 15 clusters, subdividing the 5 true cell types into further subtypes based on expression heterogeneity within each cell type.

## Conclusion

Rarity has complementary utility to existing cell type discovery methods. Rarity makes stronger assumptions about the expression patterns of distinct cell types, sacrificing sensitivity to subtle differential expression patterns to reduce the number of false positive cluster findings in order to maximise the chance of finding rare, but distinctive, cell types. We believe this construction makes Rarity particularly useful in an interactive setting in which analysts are able to manually navigate the clustering findings through Hamming Ball queries as illustrated in the examples.

Rarity has been implicitly designed for use with targeted molecular profiling technologies with data dimensions on the order of 10-40 features. We believe that in such settings, the measured molecular features will have been chosen to be cell type *markers* which makes the latent binary expression assumptions in Rarity more applicable. While, in principle, our methodological framework is general and could be extended to analyse single-cell RNA sequencing (scRNA-seq) data, considerably more care is required with the definition of “rare” cell types in high-dimensional settings. For instance, any small similar group of cells which differs from any other cell type by just a single gene could - in principle - be a candidate rare cell type. Given that a whole transcriptome analysis will yield 10,000s of genes, the possibility of large numbers of false positives is substantial.

While a number of approaches have been developed with rare cell type identification capability for scRNAseq, including RaceID3 (Grün et al. 2016), GiniClust2 (Tsoucas and Yuan 2018), GapCIust (Fa et al. 2021), CellSIUS (Wegmann et al. 2019), FiRE (Jindal et al. 2018) and scAIDE (Xie et al.2020), like in single cell cytometry, these approaches are predominantly for unsupervised discovery of larger cell populations and therefore will be sensitive to many of the issues we have discussed. Further research is required to devise a general framework for defining clustering criteria or (dis)similarity metrics that target specific cell phenotypic properties. We leave this as an open challenge to the community.

## Methods

### Conditional metrics for homogeneity and completeness

Suppose we have *C* true cell types and *K* inferred clusters. Let A be the contingency table where entry *a_ck_* denotes the number of cells corresponding to cell type *c* in cluster *k*, where the total number of cells is *N*.

Analogously to Rosenberg and Hirschberg (2007), we now define *conditional* homogeneity, completeness, and V-measure metrics which would let us “zoom in” to the rare clusters of interest as opposed to averaging across all clusters.

We first define the conditional entropies as follows:

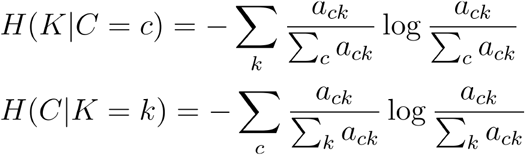

as well as the marginal entropies

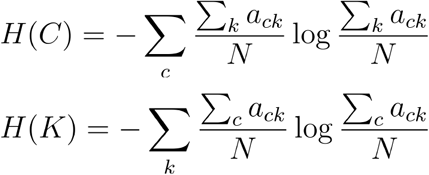

We are now interested in defining \emph{conditional} metrics to quantify the quality of the clustering w.r.t. the (potentially rare) cell type *c*. We achieve this by defining the conditional completeness

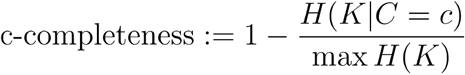

and the conditional homogeneity as

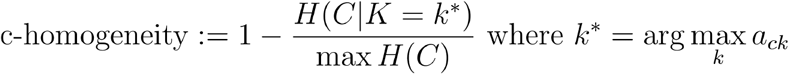

where the latter is conditional on the most likely cluster *k*^*^ for cell type *c*. Finally we define the conditional V-measure as the harmonic mean of the respective conditional completeness and homogeneity scores.

### Rarity model specification

Rarity has been designed to trade off sensitivity with respect to small cell populations and interpretability. To achieve this goal, Rarity relies on a modelling assumption that every gene is either expressed or not, and these binary states are captured via binary latent variables. Rarity implements inference for these underlying binary states. Clustering is induced by these binary signatures: cells with identical binary signatures are grouped together.

We associate every observed gene expression vector 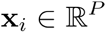 with an underlying latent variable z_i_ ∈ {0, 1}^*P*^ with binary entries *z_ig_* ~ Bernoulli(·), where *z_ig_* = 1 corresponds to gene g in cell i being expressed and *z_ig_* = 0 to being not expressed. We specify the likelihood conditional on the latent variable as follows

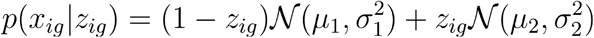

where the first component 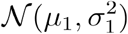 represents the distribution of markers that are “not expressed” and the second one 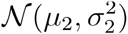 those that are “expressed”.

To make inference for this binary latent variable model tractable and scalable, we employ continuous relaxations for the binary variables in the Variational Autoencoder framework. That is, we reformulate the model as follows

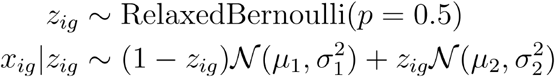

and we perform amortised variational inference with the approximate posterior *q*(**z**_i_) = RelaxedBernoulli((*f_ϕ_*(x_*i*_))) where *f_ϕ_* is an encoder neural network with shared variational parameters *ϕ*.

### Synthetic data generation

For the synthetic IMC data example, we used the following simulation scheme. We first generated the underlying binary vectors

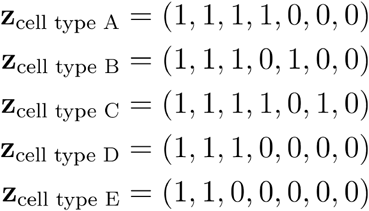

where “1” indicates that a marker is “on” and “0” that it is “off”. Conditional on these binary signatures, we then generated the observations as follows

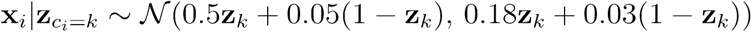

for cell type *k* {A, B, C, D, E}where the number of cells for each cell type is respectively 4000, 3000, 1000, 60, and 40.

### Sample description

Sixteen FFPE blocks of non-cancerous colon mucosa (Supplementary Table 1), were obtained from six individuals who underwent surgical resection of colorectal cancers, and subsequently reviewed by an expert pathologist. All patients provided written informed consent in accordance with approved institutional guidelines (University College London Hospital, REC Reference: 20/YH/0088; Istituto Clinico Humanitas, REC Reference: ICH-25-09).

### Staining and IMC ablation

A microtome was employed to cut one 4μm-thick section from each of the FFPE blocks from all samples. These sections were stained with a panel of 26 antibodies, targeting the main cell populations of the colon mucosa including immune, stromal and epithelial cells (Supplementary Table 2). The optimal dilution for each antibody was selected by a mucosal immunologist after reviewing the images generated from the ablation and staining of FFPE appendix sections at different concentrations (Supplementary Table 2). Before staining, slides were incubated for one-hour at 60°C, dewaxed, rehydrated and then underwent antigen retrieval. This was performed in a pressure cooker with Antigen Retrieval Reagent-Basic (R&D Systems). Then slides were blocked by incubating them in a 10% BSA (Sigma), 0.1% Tween (Sigma), and 2% Kiovig (Shire Pharmaceuticals) Superblock Blocking Buffer (Thermo Fisher) blocking solution at room temperature for two hours. The selected concentration of each antibody was added to a primary antibody mix in blocking solution and incubated overnight at 4°C. Then, the slides were washed twice in PBS and PBS-0.1% Tween and incubated for 30 minutes with the with the DNA intercalator Cell-ID™ lntercalator-lr (Fluidigm) (containing the two iridium isotopes 191lr and 193lr) 1.25 mM in a PBS solution. Subsequently the slides were washed once in PBS and once in MilliQ water and air-dried.

The Hyperion Imaging System (Fluidigm) imaging module was used to obtain a light-contrast high resolution image of approximately 4 mm^2^ of each stained slide. These images were used to select the region of interest (ROI) in each slide. 1 mm2 ROIs were selected to contain the full thickness of the colon mucosa in a longitudinal orientation, and ablated at a 1 μm/pixel resolution and 200 Hz frequency.

### IMC image processing

The ablation generated raw .txt and mcd files from which 28 images from 26 antibodies (Supplementary Table 2) and two DNA intercalators were extracted with imctools (*Imctools: Tools to Handle IMC Data n.d*.). Pixel intensities for each channel were normalised to the 99th percentile in all samples with custom R scripts and background pixels were removed by thresholding with CellProfiler2 (Kamentsky et al. 2011). A mask for the lamina propria was manually drawn for each sample using the vimentin channel as a guide, and reviewed by a mucosal immunologist.

Cells segmentation was performed first by identifying nuclei on a thresholded image derived from the multiplication of the two DNA channels. The nuclei were then used as seeds for propagation on a membrane mask derived from the sum of the E-Cadherin, CD45, CD3, CD4, CD8, CD45RO, CD27, CD68, CD34, and SMA channels. The resulting cells were then filtered according to their overlap with the lamina propria mask. Only cells overlapping the mask by more than 50% of their area were retained, resulting in a total of 40364 cells. Finally the mean pixel intensity of each marker was measured in each cell.

## Declarations

### Ethics approval and consent to participate

Patient-derived colorectal IMC data was obtained after written informed consent in accordance with approved institutional guidelines (University College London Hospital, REC Reference: 20/YH/0088; Istituto Clinico Humanitas, REC Reference: ICH-25-09).

### Availability of data and materials

We have provided the implementation of Rarity together with examples in the Github repository (https://github.com/kasparmartens/rarity). The datasets analysed during the current study are available from the Zenodo repository: (1) Breast Cancer (https://zenodo.org/record/4607374#.YgbYffXP2Lo)and (2) Colon Cancer (https://zenodo.org/record/5545882#.Yh853RPP04g).

### Authors’ contributions

FC and CY conceived the study and obtained funding. KM developed methods. MB and KM analysed data. LM and JS provided experimental support. MB, KM, FC, JS and CY interpreted results. MB, KM, FC, JS and CY wrote and edited the manuscript.

## Acknowledgements

We thank Kieran R Campbell at the University of Toronto for discussions and sharing of data and methods for Astir.

## References

Abdelaal, Tamim, Vincent van Unen, Thomas Höllt, Frits Koning, Marcel J. T. Reinders, and Ahmed Mahfouz. 2019. “Predicting Cell Populations in Single Cell Mass Cytometry Data.” Cytometry. Part A: The Journal of the International Society for Analytical Cytology 95 (7): 769–81.

Angelo, Michael, Sean C. Bendall, Rachel Finck, Matthew B. Hale, Chuck Hitzman, Alexander D. Borowsky, Richard M. Levenson, et al. 2014. “Multiplexed Ion Beam Imaging of Human Breast Tumors.” Nature Medicine 20 (4): 436–42.

Becht, Etienne, Leland McInnes, John Healy, Charles-Antoine Dutertre, Immanuel W. H. Kwok, Lai Guan Ng, Florent Ginhoux, and Evan W. Newell. 2018. “Dimensionality Reduction for Visualizing Single-Cell Data Using UMAP.” Nature Biotechnology, December, https://doi.org/10.1038/nbt.4314.

Bortolomeazzi, Michele, Lucia Montorsi, Damjan Temelkovski, Mohamed Reda Keddar, Amelia Acha-Sagredo, Michael J. Pitcher, Gianluca Basso, et al. 2022. “A SIMPLI (Single-Cell Identification from MultiPLexed Images) Approach for Spatially-Resolved Tissue Phenotyping at Single-Cell Resolution.” Nature Communications 13 (1): 781.

Bortolomeazzi, M., M. R. Keddar, and L. Montorsi. 2020. “Immunogenomic Profile of Colorectal Cancer Response to Immune Checkpoint Blockade.” bioRxiv. https://www.biorxiv.org/content/10.1101/2020.12.15.422831v1.abstract.

Cron, Andrew, Cecile Gouttefangeas, Jacob Frelinger, Lin Lin, Satwinder K. Singh, Cedrik M. Britten, Marij J. P. Welters, Sjoerd H. van der Burg, Mike West, and Cliburn Chan. 11-Jul-2013. “Hierarchical Modeling for Rare Event Detection and Cell Subset Alignment across Flow Cytometry Samples.” PLoS Computational Biology 9 (7): e1003130.

Damond, Nicolas, Stefanie Engler, Vito R. T. Zanotelli, Denis Schapiro, Clive H. Wasserfall, Irina Kusmartseva, Harry S. Nick, et al. 2019. “A Map of Human Type 1 Diabetes Progression by Imaging Mass Cytometry.” Cell Metabolism 29 (3): 755–68.e5.

Eling, Nils, Nicolas Damond, Tobias Hoch, and Bernd Bodenmiller. 2020. “Cytomapper: An R/Bioconductor Package for Visualization of Highly Multiplexed Imaging Data.” Bioinformatics 36 (24): 5706–8.

Fa, Botao, Ting Wei, Yuan Zhou, Luke johnston, Xin Yuan, Yanran Ma, Yue Zhang, and Zhangsheng Yu. 2021. “GapCIust Is a Light-Weight Approach Distinguishing Rare Cells from Voluminous Single Cell Expression Profiles.” Nature Communications 12 (1): 4197.

Gerdes, Michael J., Christopher J. Sevinsky, Anup Sood, Sudeshna Adak, Musodiq O. Bello, Alexander Bordwell, Ali Can, et al. 2013. “Highly Multiplexed Single-Cell Analysis of Formalin-Fixed, Paraffin-Embedded Cancer Tissue.” Proceedings of the National Academy of Sciences of the United States of America 110 (29): 11982–87.

Geuenich, Michael J., Jinyu Hou, Sunyun Lee, Hartland W. Jackson, and Kieran R. Campbell. 2021. “Automated Assignment of Cell Identity from Single-Cell Multiplexed Imaging and Proteomic Data.” Cell Systems. https://doi.org/10.1016/j.cels.2021.08.012.

Giesen, Charlotte, Hao A. O. Wang, Denis Schapiro, Nevena Zivanovic, Andrea jacobs, Bodo Hattendorf, Peter J. Schüffier, et al. 2014. “Highly Multiplexed Imaging of Tumor Tissues with Subcellular Resolution by Mass Cytometry.” Nature Methods 11 (4): 417–22.

Grün, Dominic, Mauro J. Muraro, Jean-Charles Boisset, Kay Wiebrands, Anna Lyubimova, Gitanjali Dharmadhikari, Maaike van den Born, et al. 2016. “De Novo Prediction of Stem Cell Identity Using Single-Cell Transcriptome Data.” Cell Stem Cell 19 (2): 266–77.

Hayday, A., and D. Gibbons. 2008. “Brokering the Peace: The Origin of Intestinal T Cells.” Mucosal Immunology.

Imctools: Tools to Handle IMC Data. n.d. Github. Accessed August 27, 2021. https://github.com/BodenmillerGroup/imctools.

Jackson, Hartland W., Jana R. Fischer, Vito R. T. Zanotelli, H. Raza Ali, Robert Mechera, Savas D. Soysal, Holger Moch, et al. 2020. “The Single-Cell Pathology Landscape of Breast Cancer.” Nature 578 (7796): 615–20.

Jindal, Aashi, Prashant Gupta, Jayadeva, and Debarka Sengupta. 2018. “Discovery of Rare Cells from Voluminous Single Cell Expression Data.” Nature Communications 9 (1): 4719.

Kamentsky, Lee, Thouis R. Jones, Adam Fraser, Mark-Anthony Bray, David J. Logan, Katherine L. Madden, Vebjorn Ljosa, Curtis Rueden, Kevin W. Eliceiri, and Anne E. Carpenter. 2011. “Improved Structure, Function and Compatibility for CellProfiler: Modular High-Throughput Image Analysis Software.” Bioinformatics 27(8): 1179–80.

Keren, Leeat, Marc Bosse, Diana Marquez, Roshan Angoshtari, Samir jain, Sushama Varma, Soo-Ryum Yang, et al. 2018. “A Structured Tumor-Immune Microenvironment in Triple Negative Breast Cancer Revealed by Multiplexed Ion Beam Imaging.” Cell 174 (6): 1373–87.e19.

Kingma, Diederik P., and Max Welling. 2014. “Auto-Encoding Variational Bayes.” Proceedings of the International Conference on Learning Representations (ICLR).

Kurd, Nadia, and Ellen A. Robey. 2014. “Unconventional Intraepithelial Gut T Cells: The TCR Says It All.” Immunity. Elsevier.

Levine, Jacob H., Erin F. Simonds, Sean C. Bendall, Kara L. Davis, El-Ad D. Amir, Michelle D. Tadmor, Oren Litvin, et al. 2015. “Data-Driven Phenotypic Dissection of AML Reveals Progenitor-like Cells That Correlate with Prognosis.” Cell 162 (1): 184–97.

Lin, Jia-Ren, Benjamin Izar, Shu Wang, Clarence Yapp, Shaolin Mei, Parin M. Shah, Sandro Santagata, and Peter K. Sorger. 2018. “Highly Multiplexed Immunofluorescence Imaging of Human Tissues and Tumors Using T-CyCIF and Conventional Optical Microscopes.” eLife 7 (July). https://doi.org/10.7554/eLife.31657.

Maaten, Laurens van der. 2008. “Visualizing Data Using T-SNE.” jmlr.org. 2008. https://www.jmlr.org/papers/volume9/vandermaaten08a/vandermaaten08a.pdf?fbclid=IwA.

Maddison, Chris J., Andriy Mnih, and Yee Whye Teh. 2017. “The Concrete Distribution: A Continuous Relaxation of Discrete Random Variables.” In International Conference on Learning Representations.

Märtens, Kaspar, Michele Bortolomeazzi, Lucia Montorsi, Jo Spencer, Francesca Ciccarelli, and Christopher Yau. 2022. “Colon Mucosa Single-Cell IMC Dataset.” Zenodo. https://doi.org/10.5281/ZENODO.6029530.

McInnes, Leland, John Healy, and james Melville. 2018. “Umap: Uniform Manifold Approximation and Projection for Dimension Reduction.” arXiv Preprint arXiv:1802.03426.

Naim, Iftekhar, Suprakash Datta, Jonathan Rebhahn, James S. Cavenaugh, Tim R. Mosmann, and Gaurav Sharma. 2014. “SWIFT-Scalable Clustering for Automated Identification of Rare Cell Populations in Large, High-Dimensional Flow Cytometry Datasets, Part 1: Algorithm Design.” Cytometry. Part A: The Journal of the International Society for Analytical Cytology 85 (5): 408–21.

Opzoomer, James W., Jessica A. Timms, Kevin Blighe, Thanos P. Mourikis, Nicolas Chapuis, Richard Bekoe, Sedigeh Kareemaghay, et al. 2021. “ImmunoCluster Provides a Computational Framework for the Nonspecialist to Profile High-Dimensional Cytometry Data.” eLife 10 (April). https://doi.org/10.7554/eLife.62915.

Paszke, Adam, Sam Gross, Francisco Massa, Adam Lerer, James Bradbury, Gregory Chanan, Trevor Killeen, et al. 2019. “PyTorch: An Imperative Style, High-Performance Deep Learning Library.” In Advances in Neural Information Processing Systems 32, edited by H. Wallach, H. Larochelle, A. Beygelzimer, F. d\textquotesingle Alché-Buc, E. Fox, and R. Garnett, 8024–35. Curran Associates, Inc.

Rezende, Danilo Jimenez, Shakir Mohamed, and Daan Wierstra. 2014. “Stochastic Backpropagation and Approximate Inference in Deep Generative Models.” arXiv Preprint arXiv:1401.4082.

Rosenberg, Andrew, and julia Hirschberg. 2007. “V-Measure: A Conditional Entropy-Based External Cluster Evaluation Measure.” In Proceedings of the 2007joint Conference on Empirical Methods in Natural Language Processing and Computational Natural Language Learning (EMNLP-CoNLL), 410–20.

Stuart, Tim, Andrew Butler, Paul Hoffman, Christoph Hafemeister, Efthymia Papalexi, William M. Mauck 3rd, Yuhan Hao, Marlon Stoeckius, Peter Smibert, and Rahul Satija. 2019. “Comprehensive Integration of Single-Cell Data.” Cell 177 (7): 1888–1902.e21.

Tsoucas, Daphne, and Guo-Cheng Yuan. 2018. “GiniClust2: A Cluster-Aware, Weighted Ensemble Clustering Method for Cell-Type Detection.” Genome Biology 19 (1): 58.

Van Gassen, Sofìe, Britt Callebaut, Mary J. Van Helden, Bart N. Lambrecht, Piet Demeester, Tom Dhaene, and Yvan Saeys. 2015. “FlowSOM: Using Self-Organizing Maps for Visualization and Interpretation of Cytometry Data.” Cytometry. Part A: The Journal of the International Society for Analytical Cytology 87 (7): 636–45.

Viney, J., T. T. MacDonald, and J. Spencer. 1990. “Gamma/delta T Cells in the Gut Epithelium.” Gut 31 (8): 841–44.

Weber, Lukas M., and Mark D. Robinson. 2016. “Comparison of Clustering Methods for High-Dimensional Single-Cell Flow and Mass Cytometry Data.” Cytometry. Part A: The Journal of the International Society for Analytical Cytology 89 (12): 1084–96.

Wegmann, Rebekka, Marilisa Neri, Sven Schuierer, Bilada Bilican, Huyen Hartkopf, Florian Nigsch, Felipa Mapa, et al. 2019. “CellSIUS Provides Sensitive and Specific Detection of Rare Cell Populations from Complex Single-Cell RNA-Seq Data.” Genome Biology 20 (1): 142.

Xie, Kaikun, Yu Huang, Feng Zeng, Zehua Liu, and Ting Chen. 2020. “scAIDE: Clustering of Large-Scale Single-Cell RNA-Seq Data Reveals Putative and Rare Cell Types.” NAR Genomics and Bioinformatics 2 (4): Iqaa082.

