## Supplementary File for "Rarity: Discovering rare cell populations from single-cell imaging data"

### Supplementary Figures

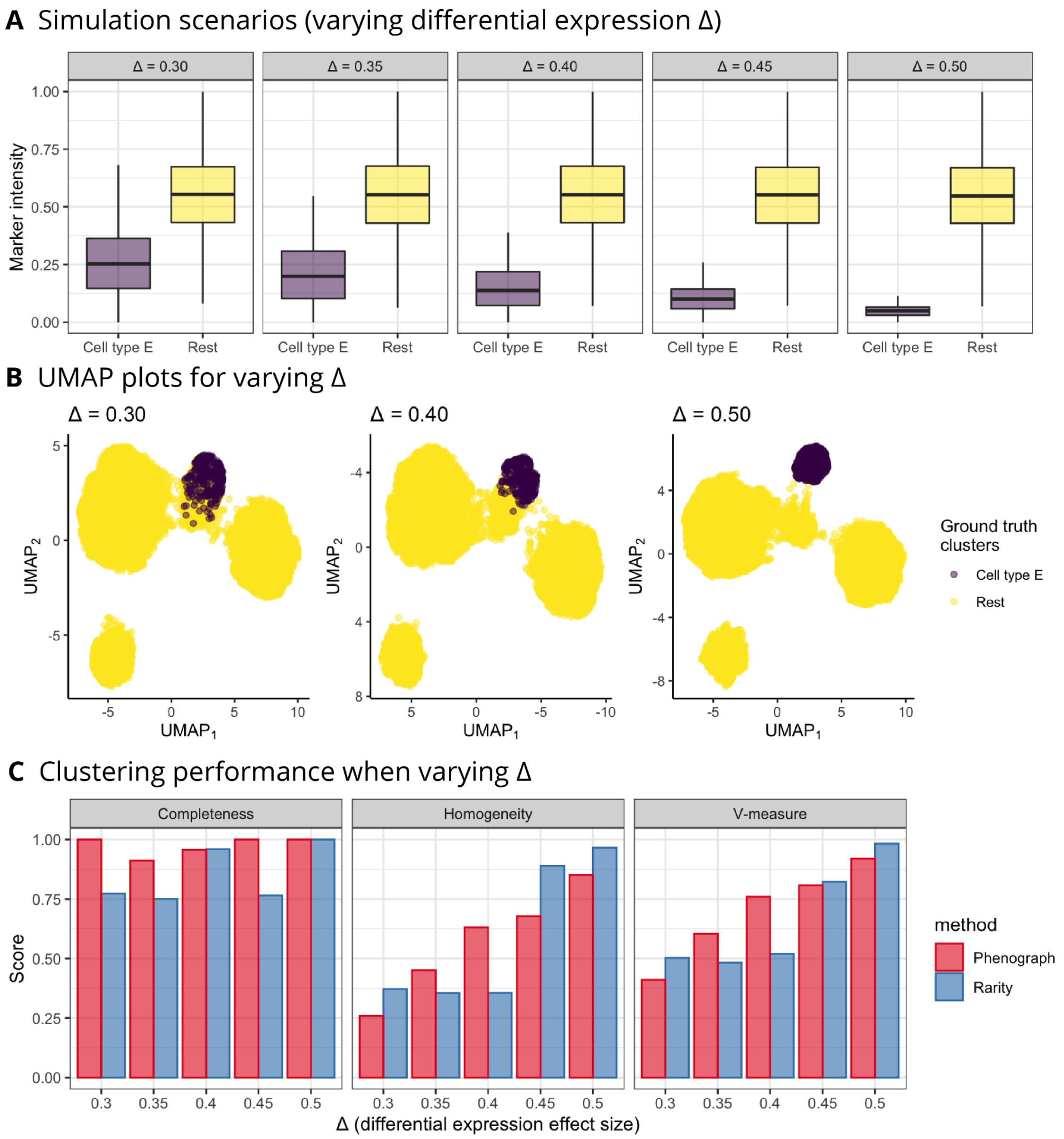


**Figure S1. Limitations of Rarity: challenging the assumption that marker genes are either "on" or "off".** Considering a synthetic data generative mechanism similar to Figures 1 and 4, here we demonstrate what happens when the discriminative marker for cell type E exhibits expression intensities decreased by amount Δ for varying levels of Δ. Specifically, for the marker shown in (A), the average intensity of all other cell types is 0.55 whereas for cell type E it is (0.55-Δ) for varying levels of Δ. The respective UMAP visualisations are shown in (B). Rarity is most successful in detecting cell type E in scenarios where Δ values are relatively large (0.50 and 0.45), as shown in panel (C), whereas its performance drops significantly for smaller values of Δ. This illustrates when Rarity's binary expression assumption breaks down - indeed for smaller Δ values the average expression intensity for cell type E starts to deviate far from zero, thus becoming challenging to distinguish for Rarity. This is in contrast to Phenograph (C) whose performance also decreases for smaller Δ values, but its performance fades more slowly as Phenograph does not take into account the absolute levels of expression.

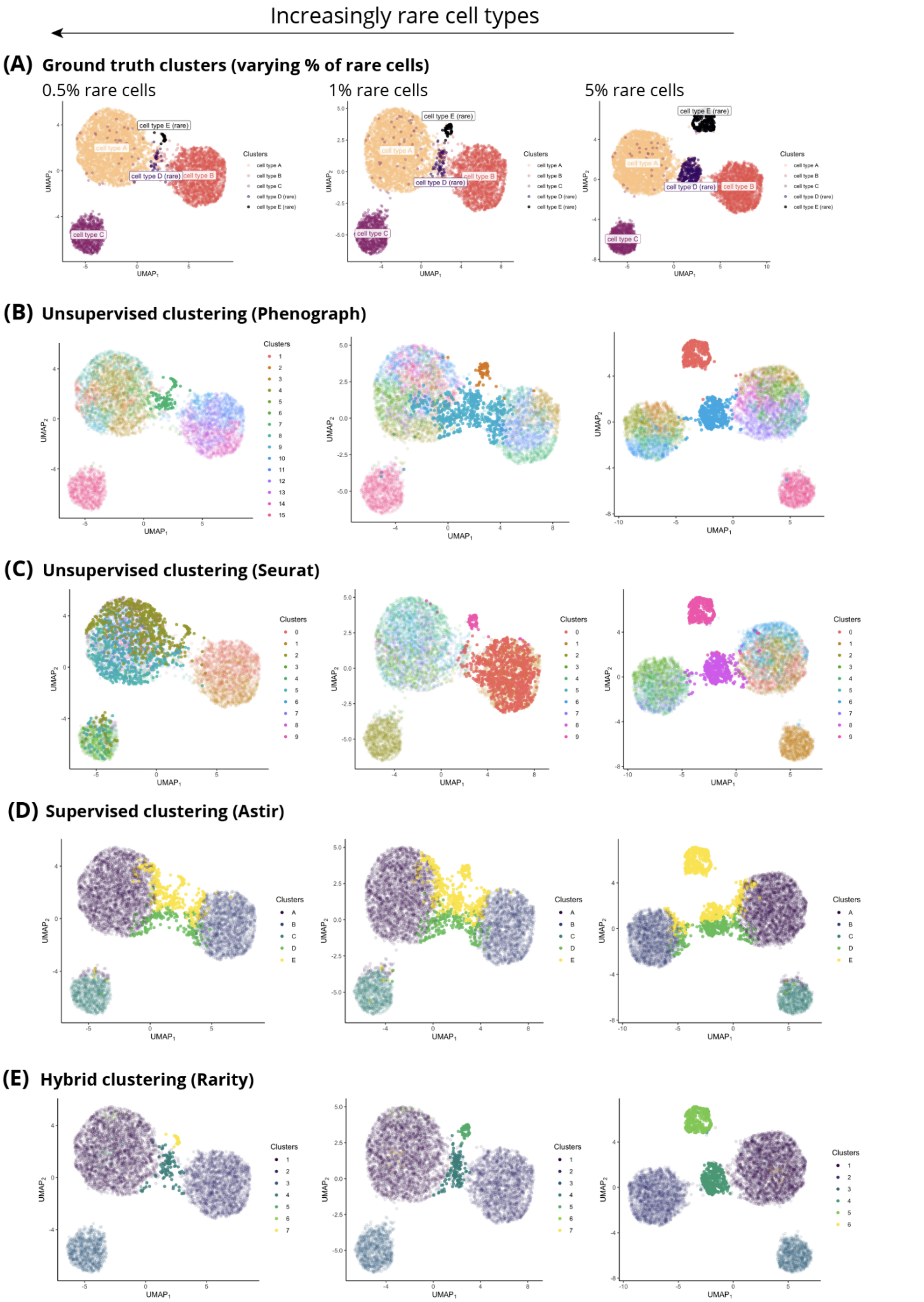


**Figure S2.** An expanded version of Figure 4, displaying (A) the ground truth in the synthetic dataset, as well as additionally displaying the clustering outcome for (B) Phenograph, (C) Seurat v3, (D) Astir, and (E) Rarity.

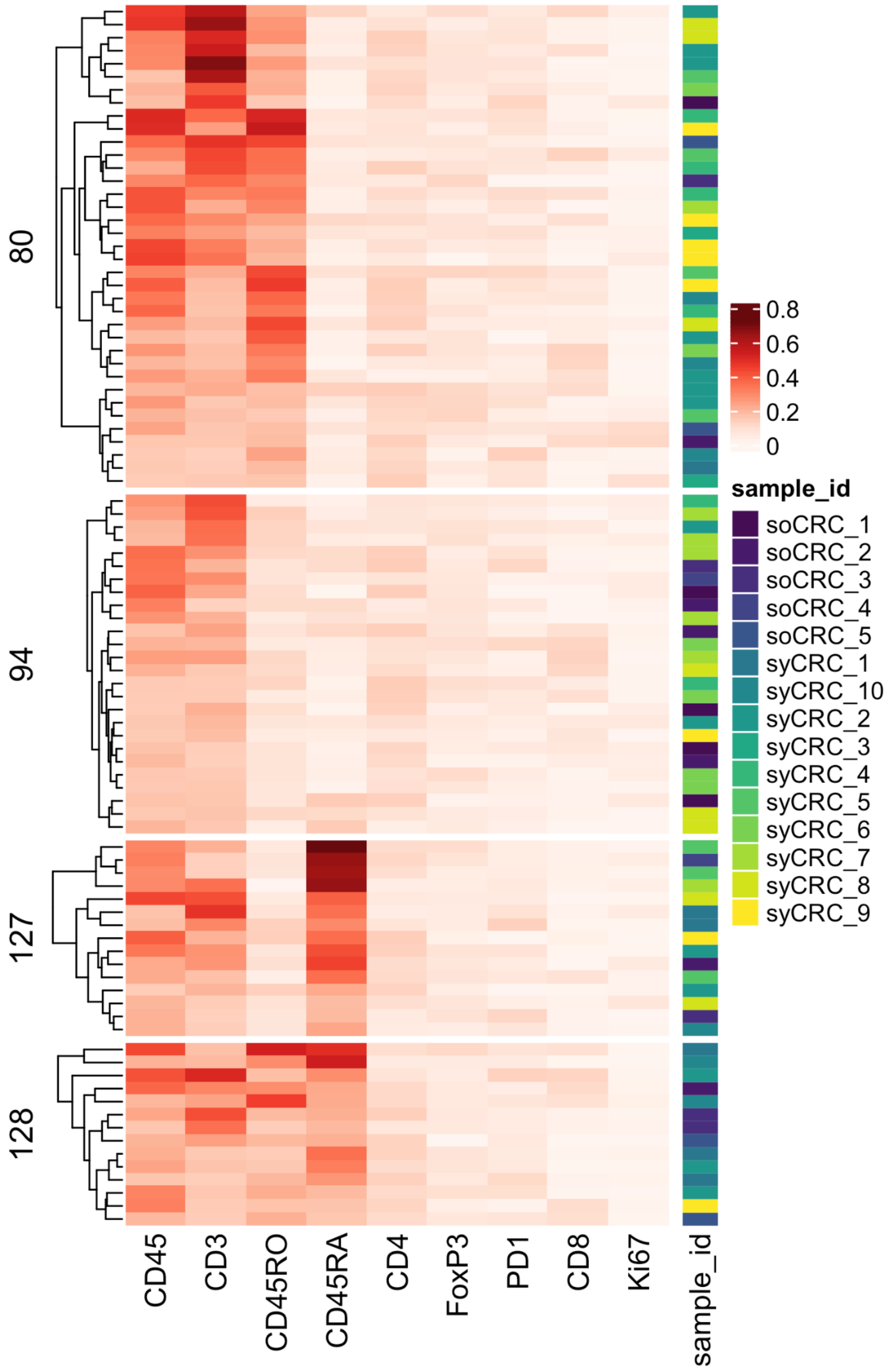


**Figure S3.** Expanded version of Figure 9D, where we show the double-negative CD4– CD8– T cell sub-groups identified by Rarity (i.e. clusters 80, 94, 127, 128), now additionally showing sample IDs (see the annotation column, coloured by sample ID). Note that these rare cells come from 15 out of 16 biological samples.

### Supplementary Tables

#### Supplementary Table 1. Samples used in the study

For each sample reported are the ID, the tissue and anatomical site of origin, the source of the sample and the figure where these data are shown. UCLH: University College London Hospital, ICH: Istituto Clinico Humanitas, NA: not available.

| **Sample ID** | **Tissue** | **Anatomical site** | **Source** | **Reference** |
|---|---|---|---|---|
| soCRC_1 | Human colon mucosa | NA | UCLH | Fig.9,10 |
| soCRC_2 | Human colon mucosa | NA | UCLH | Fig.9,10 |
| soCRC_3 | Human colon mucosa | NA | UCLH | Fig.9,10 |
| soCRC_4 | Human colon mucosa | NA | UCLH | Fig.9,10 |
| soCRC_5 | Human colon mucosa | NA | UCLH | Fig.9,10 |
| soCRC_6 | Human colon mucosa | NA | UCLH | Fig.9,10 |
| syCRC_1 | Human colon mucosa | Rectum | UCLH | Fig.9,10 |
| syCRC_2 | Human colon mucosa | Transverse colon | UCLH | Fig.9,10 |
| syCRC_3 | Human colon mucosa | Ascending colon | UCLH | Fig.9,10 |
| syCRC_4 | Human colon mucosa | Descending colon | UCLH | Fig.9,10 |
| syCRC_5 | Human colon mucosa | Rectum | UCLH | Fig.9,10 |
| syCRC_6 | Human colon mucosa | Ascending colon | ICH | Fig.9,10 |
| syCRC_7 | Human colon mucosa | Ascending colon | ICH | Fig.9,10 |
| syCRC_8 | Human colon mucosa | Descending colon | ICH | Fig.9,10 |
| syCRC_9 | Human colon mucosa | Descending colon | ICH | Fig.9,10 |
| syCRC_10 | Human colon mucosa | Ascending colon | ICH | Fig.9,10 |

#### Supplementary Table 2. Antibodies used in the study

For each antibody reported are the associated cell population, the catalogue number, the vendor, the tag, the dilution used in the staining. Data shown in figures 5, 6, and 7 were derived from a previously published breast cancer study(Jackson et al. 2020).

| Cell population | Antibody Specificity | Vendor | Catalogue Number | Metal Tag | Reference |
|---|---|---|---|---|---|
| All leukocytes | CD45 | Fluidigm | 3152016D | 152Sm | Fig.9,10 |
| B cells | CD20 | Fluidigm | 3161029D | 161Dy | Fig.9,10 |
| B cells | IgA | NovusBio | NB500-469 | 142Nd | Fig.9,10 |
| B cells | IgM | NovusBio | NBP2-34254 | 169Tm | Fig.9,10 |
| B cells / T cells | CD27 | Fluidigm | 3171024D | 171Yb | Fig.9,10 |
| T cells | CD45RA | Fluidigm | 3166028D | 166Er | Fig.9,10 |
| T cells | CD45RO | Fluidigm | 3173016D | 173Yb | Fig.9,10 |
| T cells / macrophages | CD4 | Fluidigm | 3156033D | 156Gd | Fig.9,10 |
| T cells | CD8 | Fluidigm | 3162035D | 162Dy | Fig.9,10 |
| T cells | PD1 | Fluidigm | 3165039D | 165Ho | Fig.9,10 |
| T cells | CD3 | Fluidigm | 3170019D | 170Er | Fig.9,10 |
| T cells | FOXP3 | Fluidigm | 3155016D | 155Gd | Fig.9,10 |
| Macrophages | CD68 | Fluidigm | 3159035D | 159Tb | Fig.9,10 |
| Macrophages | CD16 | Fluidigm | 3146020D | 146Nd | Fig.9,10 |
| Macrophages and dendritic cells | CD11c | Abcam | ab216655 | 175 Lu | Fig.9,10 |
| Macrophages, dendritic cells, tumour cells | PDL1 | RnD System | MAB1561 | 150Nd | Fig.9,10 |
| Endothelial cells | CD34 | Abcam | ab213058 | 164Dy | Fig.9,10 |
| Epithelial cells | Pan keratin | Fluidigm | 3148020D | 148Nd | Fig.9,10 |
| Epithelial cells | E-Cadherin | Fluidigm | 3158029D | 158Gd | Fig.9,10 |
| Basement membrane cells | Collagen type IV | NovusBio | NBP1-97716 | 176Yb | Fig.9,10 |
| Proliferating cells | Ki67 | Fluidigm | 3168022D | 168Er | Fig.9,10 |
| Stromal cells | Vimentin | Fluidigm | 3143029D | 143Nd | Fig.9,10 |
| Stromal cells | SMA | Fluidigm | 3141017D | 141Pr | Fig.9,10 |
| Various | CAMK4 | NovusBio | NBP2-37428 | 174Yb | Fig.9,10 |

| | | | | | |
|---|---|---|---|---|---|
| Various | IFNA5 | CloudClone | MAG975Hu22 | 147Sm | Fig.9,10 |
| Various | VEGFC | Abcam | ab191274 | 154Sm | Fig.9,10 |
| All Nuclei | H3 | Cell Signaling | 4499BF | In113 | Fig.5,6,7 |
| Various | H3K9me3 | Cell Signaling | 9733BF | La139 | Fig.5,6,7 |
| Basal epithelium | CK5 | Abcam | Custom | Pr141 | Fig.5,6,7 |
| Stromal cells | Fibronectin | BD Biosciences | 610078 | Nd142 | Fig.5,6,7 |
| Luminal epithelium | CK19 | Dev Studies Hybridoma Bank Troma-III | 37815 | Nd143 | Fig.5,6,7 |
| Luminal epithelium | CK8/18 | Cell Signaling | 4546BF | Nd144 | Fig.5,6,7 |
| Various | Twist | Millipore | ABD29 | Nd145 | Fig.5,6,7 |
| Macrophages | CD68 | E-Bioscience | 14-0688-82 | Nd146 | Fig.5,6,7 |
| Basal epithelium | KRT14 | Thermo Fischer | PA5-16722 | Sm147 | Fig.5,6,7 |
| Stromal cells | SMA | Abcam | ab7817 | Nd148 | Fig.5,6,7 |
| Stromal cells | Vimentin | Cell Signaling | 5741BF | Sm149 | Fig.5,6,7 |
| Various | c-Myc | Biolegend | 626802 | Nd150 | Fig.5,6,7 |
| Her2 cancer | HER2 | BD Biosciences | 554299 | Eu151 | Fig.5,6,7 |
| T Cells | CD3ε | Cell Signaling | 85061 | Sm152 | Fig.5,6,7 |
| All Nuclei | H3 | Biolegend | 641002 | Eu153 | Fig.5,6,7 |
| Various | Slug | R&D Systems | Custom | Gd155 | Fig.5,6,7 |
| ERα+ cancer | ERα | Epitomics | AC-0015EU | Gd156 | Fig.5,6,7 |

| | | | | | |
|---|---|---|---|---|---|
| PR+ cancer | PR A/B | Spring Bioscience | M3024 C | Gd158 | Fig.5,6,7 |
| PR+ cancer | PR A/B | Epitomics | AC-0028EU | Gd158 | Fig.5,6,7 |
| All cells | p53 | Cell Signaling | 2527BF | Tb159 | Fig.5,6,7 |
| Various | CD44 | R&D Systems | AF3660 | Gd160 | Fig.5,6,7 |
| All leukocytes | CD45 | E-Bioscience | 14-9457-82 | Dy162 | Fig.5,6,7 |
| Epithelilal cells | GATA3 | BD Bioseeinces | 558686 | Dy163 | Fig.5,6,7 |
| B cells | CD20 | E-Bioscience | 14-0202-82 | Dy164 | Fig.5,6,7 |
| Various | CA9 | R&D Systems | AF2188 | Er166 | Fig.5,6,7 |
| Epithelilal cells | E-Cadherin/P-Cadherin | BD Biosciences | 610182 | Er167 | Fig.5,6,7 |
| Proliferating cells | Ki67 | Cell Signaling | 9449BF | Er168 | Fig.5,6,7 |
| EGFR+ cancer | EGFR | Cell Signaling | 4267BF | Tm169 | Fig.5,6,7 |
| Various | p-S6 | Cell Signaling | 4858BF | Yb170 | Fig.5,6,7 |
| Various | vWF | Millipore | AB7356 | Yb172 | Fig.5,6,7 |
| Endothelial cells | CD31 | Novus Biologicals | NB600-562 | Yb172 | Fig.5,6,7 |
| Various | p-mTOR | Cell Signaling | 2976 | Yb173 | Fig.5,6,7 |
| Luminal epithelium | CK7 | Biosciences | 550507 | Yb174 | Fig.5,6,7 |
| Epithelilal cells | Pan CK | MAB1612 | 2341224 | Lu175 | Fig.5,6,7 |
| Epithelilal cells | Pan CK | MAB1611 | 2607604 | Lu175 | Fig.5,6,7 |
| Apoptosis | cleaved PARP | BD Biosciences | 552596 | Yb176 | Fig.5,6,7 |

| | | | | | |
|---|---|---|---|---|---|
| Apoptosis | Cleaved Caspase3 | BD Biosciences | 559565 | Yb176 | Fig.5,6,7 |

#### Supplementary Table 3. Cell type definition for epithelial luminal cells (for Figure 6)

We defined epithelial luminal cells as those expressing the following markers:

- DNA1, DNA2
- E-cadherin
- Pan-Cytokeratin
- Cytokeratin 7
- Cytokeratin 8/18
- Cytokeratin 19

whereas not expressing the following markers

- Vimentin
- vWF
- SMA
- CD45
- CD3
- CD20
- CD68

#### Supplementary Table 4. Cell type definitions for colon mucosa data (used in Figure 9A)

| | CD45 | CD3 | CD68 | CD20 | IgA | CD11c | CD34 | SMA | eCadherin | CollagenIV |
|---|---|---|---|---|---|---|---|---|---|---|
| T cells | 1 | 1 | 0 | 0 | | | | | | |
| B cells | 1 | 0 | 0 | One of CD20/IgA | | | | | | |
| Macrophages | 1 | 0 | 1 | 0 | | | | | | |
| Dendritic cells | 0 | | | | | 1 | | 0 | | |
| Epithelial | 0 | | | | | | | | 1 | |
| Endothelial | 0 | | | | | | 1 | | | |
| Connective | 0 | | | | | | | 1 | | 1 |
